# Chemogenetic analysis of how receptors for short chain fatty acids regulate the gut-brain axis

**DOI:** 10.1101/2020.01.11.902726

**Authors:** Natasja Barki, Daniele Bolognini, Ulf Börjesson, Laura Jenkins, John Riddell, David I. Hughes, Trond Ulven, Brian D. Hudson, Elisabeth Rexen Ulven, Niek Dekker, Andrew B. Tobin, Graeme Milligan

## Abstract

The gut-brain axis allows bi-directional communication between the enteric and central nervous systems. Short chain fatty acids (SCFAs) generated by the gut microbiota are important regulators of this interface. However, defining mechanisms by which SCFAs do so has been challenging because, amongst various roles, they co-activate both of a pair of closely related and poorly characterized G protein-coupled receptors, FFA2 and FFA3. Designer Receptors Exclusively Activated by Designer Drugs (DREADDs) can provide an important approach in defining receptor-specific functions. By screening a library of carboxylate-containing small molecules we identified 4-methoxy-3-methyl-benzoic acid (MOMBA) as a specific agonist of a DREADD variant of FFA2 which is not activated by SCFAs. Using mice engineered to replace FFA2 with this FFA2-DREADD, whilst retaining FFA3 expression, combinations of MOMBA and the now FFA3 receptor selective SCFAs defined key, but distinct, roles of FFA2 and FFA3 in each of gut transit time, secretion of entero-endocrine hormones, and communication from the gut to each of autonomic and somatic sensory ganglion cells and the spinal cord. These studies map mechanisms and signalling pathways by which each of FFA2 and FFA3 act to link the gut and the brain and provide both animal models and novel tool compounds to further explore this interface.

## Introduction

The gut-brain axis allows bi-directional communication between the enteric and central nervous systems. Growing evidence highlights the role that the intestinal microbiota may play in such interactions (1) and in the development of disease (2). The microbiota generates a wide array of metabolites that can modulate host cells and their functions (3–4). Among these short chain fatty acids (SCFAs), particularly acetate (C2) and propionate (C3), are generated in prodigious amounts by fermentation of fibre and other non-digestible carbohydrates in the lower gut. The SCFAs play central roles in homeostasis at the interface between metabolism and immunity (5–6) and also within the gut-brain axis (7–8). Although a range of their key effects are believed to be produced by activation of G protein-coupled receptors (GPCRs), both locally in the lower gut and following uptake into the systemic circulation, which of these roles are generated directly by individual receptors and via which signalling pathways remains uncertain.

Over the course of evolution genes encoding many ancestral forms of GPCRs have multiplied and diversified to provide enhanced flexibility of signalling and integration of responses to either the same or closely related ligands. An example of this is within the family of receptors that are activated by the binding of free fatty acids. In human, genes encoding the FFA1, FFA2 and FFA3 receptors cluster at chromosomal location 19q13.1 and in mouse, in a similar manner, at chromosomal location 7 A3 (9). Each receptor is closely related and whilst FFA1 is activated by saturated and unsaturated fatty acids of chain length C10 and above, both FFA2 and FFA3 are activated instead by SCFAs (9–10). There is marked overlap between the potency of SCFAs at FFA2 and FFA3 but variation between the optimal chain length required to activate mouse versus human FFA2 and FFA3 (11). However, SCFAs also have multiple non-GPCR-mediated effects in the body and this poses additional challenges in using these molecules to define effects as being mediated by either FFA2 or FFA3 (9–10). There is a paucity of highly selective pharmacological tool compounds for FFA2 and FFA3 (12–13), particularly that are equally effective at human and rodent orthologs of these receptors. To attempt to provide selective pharmacology, in early studies Schmidt et al., (14) explored the selectivity between FFA2 and FFA3 of a number of small carboxylic acids. Although a degree of selectivity was observed, and some of these compounds have subsequently been used as modestly selective FFA2 or FFA3 ligands (15–17), they are far from satisfactory.

To overcome these issues, and to define unambiguously specific roles of FFA2 and FFA3 in the actions of SCFAs at various levels within the gut-brain axis, herein we have applied an integrated chemogenetic approach. Firstly, we have extended the use of a transgenic knock-in mouse line in which we expressed a Designer Receptor Exclusively Activated by Designer Drugs (DREADD) derived from human (h) FFA2 (18) in place of endogenous mouse FFA2 (19). This variant form does not respond to C2 or C3 and, therefore, both *in vivo* and in cells and tissues derived from these animals responses to C2 or C3 cannot reflect activation of hFFA2-DREADD. However, expression of FFA3 is unaffected in these animals (19) and, therefore, effects of the SCFAs may instead be mediated by FFA3. In previous studies we identified that 4-hexadienoic acid (sorbic acid) acts as an agonist at hFFA2-DREADD but is without activity at various wild type orthologs of FFA2 and of FFA3 (18–19). As such, effects of sorbate in the transgenic mice are potentially mediated by hFFA2-DREADD but not by FFA3 (19).

Although we are unaware of any noted off-target effects of sorbate, and it has extremely low toxicity (20), it is widely used as a food preservative (21). Because of its potential anti-bacterial and anti-fungal effects, which might affect longer-term studies in which we would provide sorbate in the drinking water of animals, we wish to identify other, chemically distinct, hFFA2-DREADD agonist ligands. A richer chemical toolbox of activators is, moreover, always desirable for selectivity and functional studies for any pharmacological or therapeutic target. By screening libraries of small molecules with similarity to sorbate we identified a series of 4-methoxy-benzoic acid derivatives that also act as highly selective activators of hFFA2-DREADD. From these we selected 4-methoxy-3-methyl-benzoic acid (MOMBA) for the studies reported herein.

Using the unique FFA2-DREADD transgenic mice and combinations of MOMBA and C3, alongside a novel FFA3 selective activator (TUG-1907) we recently identified (22) we have dissected the contributions of FFA2 and FFA3 to functions of SCFAs at levels ranging from gut transit to the activation of dorsal root- and nodose-ganglion cells and the spinal cord. We show that combinations of these two receptors transduce different effects of SCFAs within these pathways and via different G protein-mediated pathways and by so doing control the microbiota-gut-brain axis.

## Results

### Identification of 4-methoxy-benzoic acid derivatives as novel hFFA2-DREADD agonists

We have previously shown that sorbate is a moderately potent, selective and effective agonist of hFFA2-DREADD that lacks agonist action at each of human and mouse FFA2 and both human and mouse FFA3 (18–19). However, for pharmacological studies, particularly when exploring native cells and tissues, use of more than a single receptor-activating ligand is important to help define with certainty ‘on-target’ and functionally relevant responses. We, therefore, conducted a screen for novel agonists at hFFA2-DREADD using initially a receptor-β-arrestin-2 interaction assay. Here, following transient transfection of HEK293 cells to express both hFFA2-DREADD tagged at the intracellular C-terminal tail with enhanced Yellow Fluorescent Protein (hFFA2-DREADD-eYFP) and β-arrestin-2-*Renilla*-luciferase, induced proximity of eYFP and *Renilla*-luciferase allows bioluminescence resonance energy transfer (BRET) that reflects agonist-promoted interactions between the hFFA2-DREADD receptor and the arrestin (19). Using sorbate as a positive control, BRET signal was enhanced in a concentration-dependent manner with pEC_50_ = 3.89 +/- 0.04 (n = 3) **Figure 1A**). Using this assay we initially screened, at 100 μM, more than 1200 small molecules selected to have some structural similarity with sorbate and designed/collected to have good physicochemical properties and a lack of known chemical liabilities (**Supplementary Figure 1A**). This provided a robust 96 well microtitre plate-based assay with calculated Z’ (23) routinely > 0.6 (**Supplementary Figure 1B)** and greater than 5-fold signal to background (**Supplementary Figure 1C**). Reconfirmation screens of potential hits, also conducted at 100 μM, provided positives, including compounds 565 and 1184 (**Figure 1B**). Deconvolution indicated that compound 1184 was, in fact, sorbic acid, confirming the capacity of the screen to identify effectively a previously characterized active, whilst compound 565 was 4-methoxy-3-methyl-benzoic acid (MOMBA) (**Figure 1B**). Potential activity of a further 320 distinct compounds, selected now on relatedness to the hits from the initial screen, was again assessed initially at a single concentration (100 μM) in 96 well format (**Figure 1C**). This resulted in the identification of further compounds including a number closely related to MOMBA, such as 4-methoxy-3-chloro-benzoic acid (compound 132) and 4-methoxy-3-hydroxy-benzoic acid (compound 235) as actives (**Figure 1C**). A number of these were at least as potent as sorbate, with the best displaying some 3-fold higher potency in this assay (**Figure 1D**). Based on the common 4-methoxy-3-X-benzoic acid scaffold of MOMBA and compounds 132 and 235, MOMBA was selected for more detailed studies and purchase of MOMBA from a separate source confirmed activity of this chemical.

**Figure 1.**
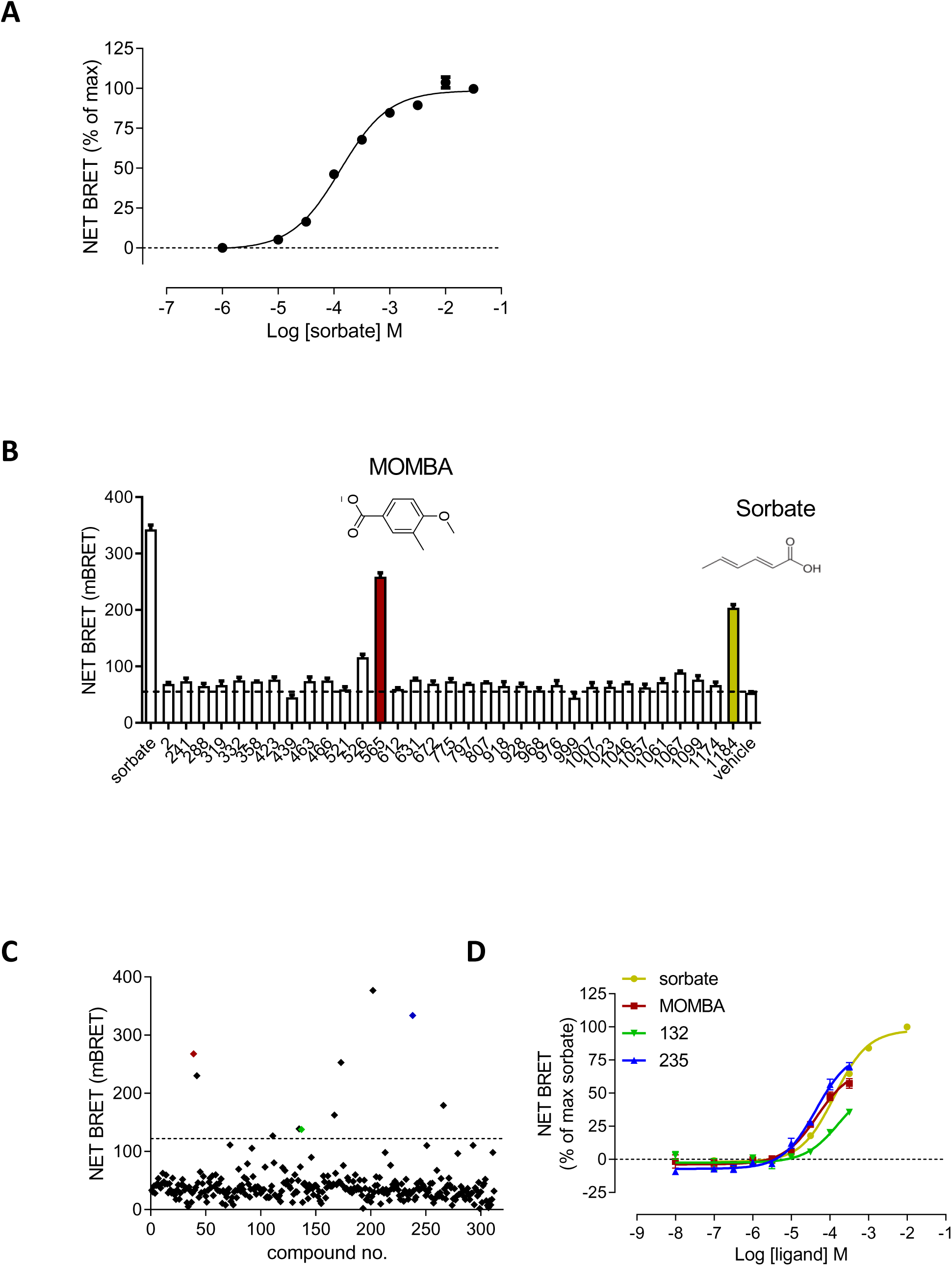
Screening for novel agonists of hFFA2-DREADD identifies MOMBA. A. HEK293 cells were transiently transfected to express both hFFA2-DREADD-eYFP and β-arrestin-2-*Renilla*-luciferase. Addition of sorbate promoted interactions between these proteins in a concentration-dependent manner. Data are means +/- S.E.M. of triplicates from a single experiment, representative of 6. B. A subset of the positives from the initial screen (see **Supplementary Figure 1A**) were re-tested at 100 μM. Compounds 565 (MOMBA) (**red**) and 1184 (sorbate) (**yellow**) are highlighted. Dotted line indicates basal signal. Data are from a single experiment with results plotted as mean +/- S.E.M. of triplicate assays. C. A further 320 compounds selected on similarity to hits from **Supplementary Figure 1A** were selected and screened at 100 μM in single point assays: As well as MOMBA (**red**) two of these were 4-methoxy-3-chloro-benzoic acid (compound 132) (**green**) and 4-methoxy-3-hydroxy-benzoic acid (compound 235) (**blue**). Dotted line indicates selection cut-off. Data are from a single experiment. D. Concentration-dependence of selected hits from **C** to activate hFFA2-DREADD is displayed. MOMBA (**red**), compound 132 (**green**) and compound 235 (**blue**). Sorbate (**yellow**) is shown as reference.

### MOMBA is a highly selective agonist of hFFA2-DREADD

To be useful as hFFA2-DREADD specific agonists compounds should not activate either wild type human or mouse FFA2. For MOMBA this was assessed initially for wild type hFFA2 using an equivalent BRET-based receptor-β-arrestin-2 interaction assay to that described above for hFFA2-DREADD. Following transient co-expression of hFFA2-eYFP and β-arrestin-2-*Renilla*-luciferase in HEK293 cells, whilst the SCFA propionate (C3) was an effective agonist with pEC_50_ = 3.46 +/- 0.01 (n = 3) MOMBA was without detectable activity (**Supplementary Figure 2A**). For reasons that remain undefined mouse FFA2 did not allow development of an equivalent, suitably robust receptor-β-arrestin-2 interaction assay. As such, because orthologs of FFA2 can activate G_i_-family G proteins (24–25) we turned to inhibition of cAMP assays. In Flp-In T-REx 293 cells stably harboring hFFA2-eYFP, but in which expression of the receptor construct was not induced, none of C3, sorbate or MOMBA was able to reduce levels of forskolin-stimulated cAMP (**not shown**). In contrast, after hFFA2-eYFP expression was induced in these cells by addition of doxycycline, C3 inhibited forskolin-amplified cAMP levels, but neither sorbate nor MOMBA did so (**Supplementary Figure 2B**). Equivalent results were produced in Flp-In T-REx 293 cells induced to express mouse FFA2-eYFP (**Supplementary Figure 2B**), where again only C3 but not either sorbate or MOMBA was able to inhibit forskolin amplified cAMP levels. The closely related receptor FFA3 is also activated by SCFAs including C3 (24–25). It was, therefore, vital for subsequent studies that MOMBA also lacked activity at this receptor. Using equivalent Flp-In T-REx 293 cell lines able to induce expression of either human or murine forms of FFA3-eYFP, whilst C3 was once again able to inhibit cAMP levels via both orthologs neither sorbate nor MOMBA was effective (**Supplementary Figure 2C**).

The ability of MOMBA to inhibit forskolin-stimulated cAMP in cells expressing the hFFA2-DREADD receptor was concentration-dependent and MOMBA was more potent (pEC_50_ = 5.24 +/- 0.16, mean +/- S.E.M. n = 3) than sorbate (pEC_50_ = 4.77 +/- 0.15, mean +/- S.E.M. n = 3) in so doing (**Supplementary Figure 3A**). FFA2 is also able to interact with and activate G_q_-family G proteins as well as G_i_-family members (19, 25–26). We established, therefore, that although C3 was unable to promote inositol monophosphate generation in Flp-In T-REx 293 cells induced to express hFFA2-DREADD (19), MOBMA again did so in a concentration-dependent manner and with both potency and efficacy at least equivalent to sorbate (**Supplementary Figure 3B**). These characteristics were re-iterated in measures of ligand-regulated binding of [^35^S]GTPγS performed on membranes from Flp-In T-REx 293 cells induced to express hFFA2-DREADD (**Supplementary Figure 3C**). A key reason for using human FFA2 as the basis for the DREADD construct was that available antagonist ligands are able to block human FFA2, but not the mouse ortholog (26–27). The effect of an EC_80_ concentration of MOMBA to stimulate [^35^S]GTPγS binding was fully inhibited by increasing concentrations of both of the structurally distinct, human FFA2 ortholog specific, antagonists 4-[[[(2*R*)-1-(benzo[*b*]thien-3-ylcarbonyl)-2-methyl-2-azetidinyl]carbonyl][(3-chlorophenyl)methyl]amino]butanoic acid (GLPG0974) (pIC_50_ = 7.58 +/- 0.07, mean +/- S.E.M. n = 3) and (((*S*)-3-(2-(3-chlorophenyl)acetamido)-4-(4-(trifluoromethyl)phenyl)butanoic acid) (CATPB) (pIC_50_ = 7.63 +/- 0.08, mean +/- S.E.M. n = 3) (**Supplementary Figure 3D**). This is consistent with, as for sorbate (19), MOMBA likely binding in the orthosteric pocket of hFFA2-DREADD. This was confirmed as MOMBA lacked activity at an Arg^180^Ala mutant of hFFA2-DREADD (**Supplementary Figure 3E**). Mutation of Arg^180^ in FFA2 is known to eliminate binding and activity of carboxylate-containing orthosteric agonists of FFA2 (25).

### Selective pharmacology of activation of hFFA2-DREADD-HA and mouse FFA3

We recently generated a ‘knock-in’ transgenic mouse line in which mouse FFA2 is replaced with a humanized sequence able to encode a C-terminally HA-epitope-tagged form of hFFA2-DREADD (19). Driven by the endogenous *ffar2* gene promoter regions this line expresses hFFA2-DREADD-HA in the same tissues as mouse FFA2 in wild type animals, and to similar levels (19). The experiments described above defined MOMBA as a hFFA2-DREADD specific agonist, with improved potency compared to sorbate, but with no activity at FFA3. However, because C3 effectively activates mouse FFA3 but not hFFA2-DREADD (**Supplementary Figure 2**) (18–19), this defined selective pharmacological agonists for each of these receptors in tissues of the hFFA2-DREADD-HA knock-in mouse line that would potentially allow assessment of the contributions of each of FFA2 and FFA3 at various levels within the microbiota-gut-brain axis. It has already been established that both FFA2 and FFA3 are expressed in subsets of entero-endocrine cells of the gut (19, 28), and that FFA3 is expressed in a range of autonomic and somatic sensory ganglia (28–30) however their roles in connectivity to the gut are undefined.

### FFA2 but not FFA3 activation promotes gut transit time

One of the more uncertain aspects of mechanisms behind local effects of SCFA in the gut following their production has been on gut transit time. For example, both (31) and (32) observed SCFAs to reduce gut transit. By contrast Bolognini et al., (19) showed that in wild type C57BL/6 mice provision of C3 in the drinking water resulted in enhanced gut transit. This could reflect activation of FFA2, FFA3, a combination of the two GPCRs, or neither of them. To ensure that MOMBA could act effectively *in vivo* following oral administration we initially provided MOMBA (15 mM) in the drinking water of hFFA2-DREADD-HA expressing mice. This significantly increased (P < 0.01, n = 5) gut transit in these animals compared to water alone (**Figure 2A**). Importantly, subsequent removal of MOMBA from the drinking water restored transit time in the same individual mice (P < 0.01) to control levels (**Figure 2A**). This clearly was an on-target, hFFA2-DREADD-mediated effect of MOMBA because provision of MOMBA in the drinking water of either wild type C57BL/6 mice (**Figure 2B**) or ‘CRE-MINUS’ mice (19), in which neither mouse FFA2 nor hFFA2-DREADD-HA is expressed (**Figure 2C**), was without effect on gut transit time. We have previously shown that provision of C3 (150 mM) in the drinking water of C57BL/6 mice also increases gut transit time (19), but in concert with the effects of MOMBA reported here and that C3 has no effect in mice lacking expression of FFA2 or in which hFFA2-DREADD-HA is expressed instead of mouse FFA2 (19), this also clearly reflects activation of FFA2 in wild type mice, rather than FFA3 or a non-receptor mediated effect.

**Figure 2.**
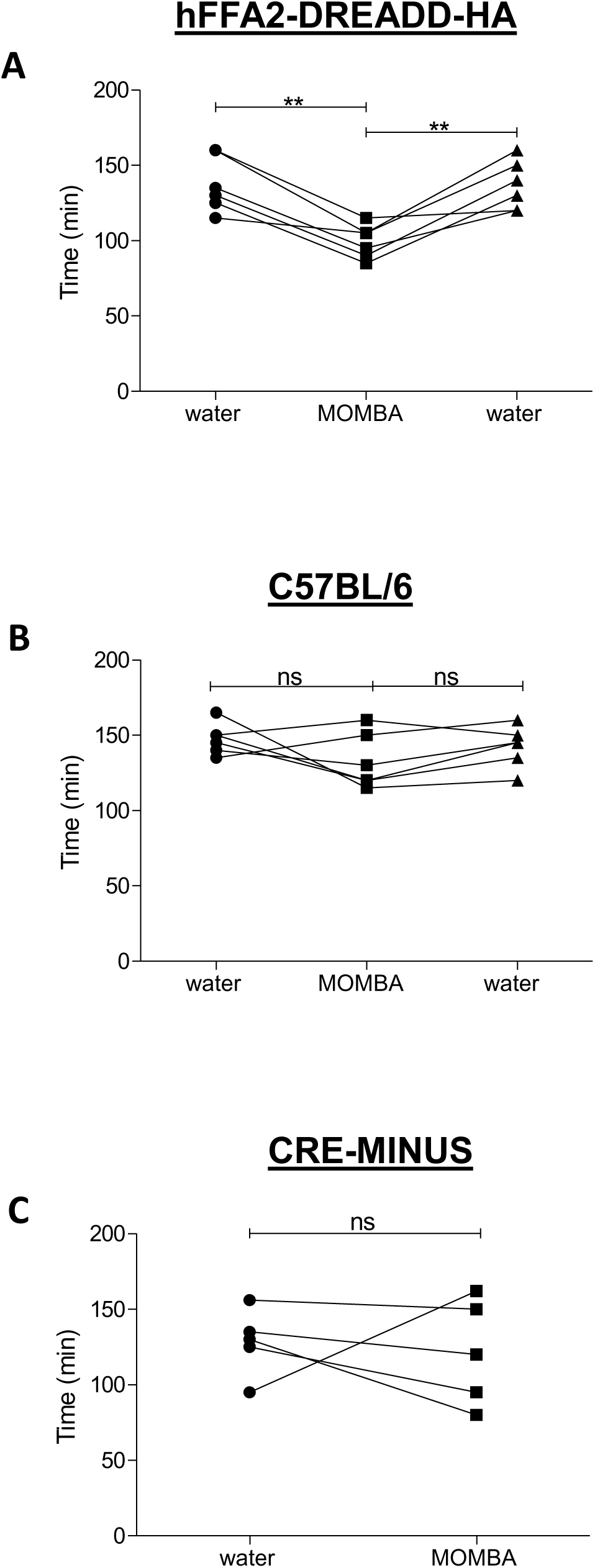
MOMBA promotes increased gut transit via hFFA2-DREADD-HA. Male hFFA2-DREADD-HA (**A**), wild type C57BL/6 (**B**) and CRE-MINUS (**C**) mice were acclimatized for 7 days with free access to drinking water. Individual animals were then gavaged with carmine red and total GI transit time measured. Following the initial transit studies mice were provided with MOMBA (15 mM) in the drinking water. After a further 7 days GI transit of all mice was again measured. In **A** and **B** MOMBA was then removed and the mice were again provided with water followed by a further gavage with carmine red 7 days later. Data are for individual animals (** P < 0.01, ns = not significant).

### Colonic release of GLP-1 is mediated specifically by FFA2

Both FFA2 and FFA3 are expressed by various enteroendocrine cells throughout the small and large intestine (28, 33–34), and activation of both these GPCRs by SCFAs has been implicated in the release of incretins such as glucagon-like peptide 1 (GLP-1) (28, see 33-34 for review). We, therefore, next isolated colonic crypt-containing preparations from the hFFA2-DREADD-HA transgenic mice. As demonstrated previously (19) addition of the cAMP-phosphodiesterase inhibitor isobutylmethylxanthine (IBMX) resulted in a large increase in release of GLP-1 from these (**Supplementary Figure 4A**). MOBMA promoted release of GLP-1 in a concentration-dependent manner, in which a statistically significant (P < 0.05) effect could be detected by exposure to 1 μM MOMBA (**Supplementary Figure 4A**). Moreover, at 100 μM MOMBA was as effective as 1 mM sorbate (**Supplementary Figure 4A**), consistent with the higher potency of MOMBA measured in the 293 cell line-based studies (**Supplementary Figure 3**) and with MOMBA being at least as efficacious as sorbate in native cells and tissues as well as in transfected cell lines. By contrast C3 (10 mM) was unable to produce such an effect, indicating GLP-1 release from these preparations to be a selective attribute of activation of hFFA2-DREADD-HA and not to require activation of FFA3. To examine this further, and in a more physiologically intact preparation, the colon was dissected from hFFA2-DREADD-HA mice and mounted in the chamber of an organ bath in which ligands could be perfused through the tissue. Once baseline GLP-1 release was established MOMBA was introduced into the buffer. This caused a rapid and sustained increase in release of GLP-1 (**Supplementary Figure 4B**). We have shown previously that the mutations introduced into hFFA2 to generate the DREADD variant receptor do not interfere with binding of the hFFA2 ortholog specific antagonist CATPB (19) and as shown in **Supplementary Figure 3D** CATPB effectively antagonises effects of MOMBA in 293-derived cells transfected to express hFFA2-DREADD. When colonic tissue from hFFA2-DREADD-HA mice was exposed to a combination of CATPB and MOMBA no release of GLP-1 above basal levels could be detected (**Supplementary Figure 4B**), further confirming the specific role of hFFA2-DREADD. A ‘CRE-MINUS’ strain of these mice harbors hFFA2-DREADD-HA under the control of *ffar2* gene promoter regions but neither the hFFA2-DREADD-HA construct nor mouse FFA2 is expressed (19). These animals are thus functionally akin to a FFA2 knock-out line (19). In colonic tissue from these CRE-MINUS animals MOMBA was unable to promote release of GLP-1 (**Supplementary Figure 4C**), further confirming this endpoint to reflect activation of hFFA2-DREADD-HA rather than any potential off-target, non-receptor, mediated effect of MOMBA.

### Colonic release of PYY is also mediated specifically by FFA2

Peptide YY (PYY) is expressed in similar subsets of entero-endocrine ‘L-cells’ as GLP-1 (28, 34) and perfusion of colonic tubes isolated from hFFA2-DREADD-HA mice with buffer containing MOMBA also resulted in a rapid, markedly enhanced (P < 0.01), but in this case transient, release of PYY (**Figure 3A**). This effect of MOMBA was not observed in the presence of CATPB (**Figure 3A**) and was also not produced in equivalent preparations generated from the CRE-MINUS animals (**Figure 3B**). A distinct feature of the hFFA2-DREADD-HA expressing mice is that the appended C-terminal HA epitope tag allows exquisite immunochemical detection of cells expressing the receptor (19). Consistent with the MOMBA-induced release of PYY, co-staining of colonic sections from the hFFA2-DREADD-HA mice with both anti-HA and anti-PYY antibodies identified a subset of cells that as well as being positive for PYY also expressed hFFA2-DREADD-HA (**Figure 3C**). Such co-localized staining was lacking in tissue sections isolated from CRE-MINUS animals although identification of PYY expressing cells was equivalent (**Figure 3C**).

**Figure 3.**
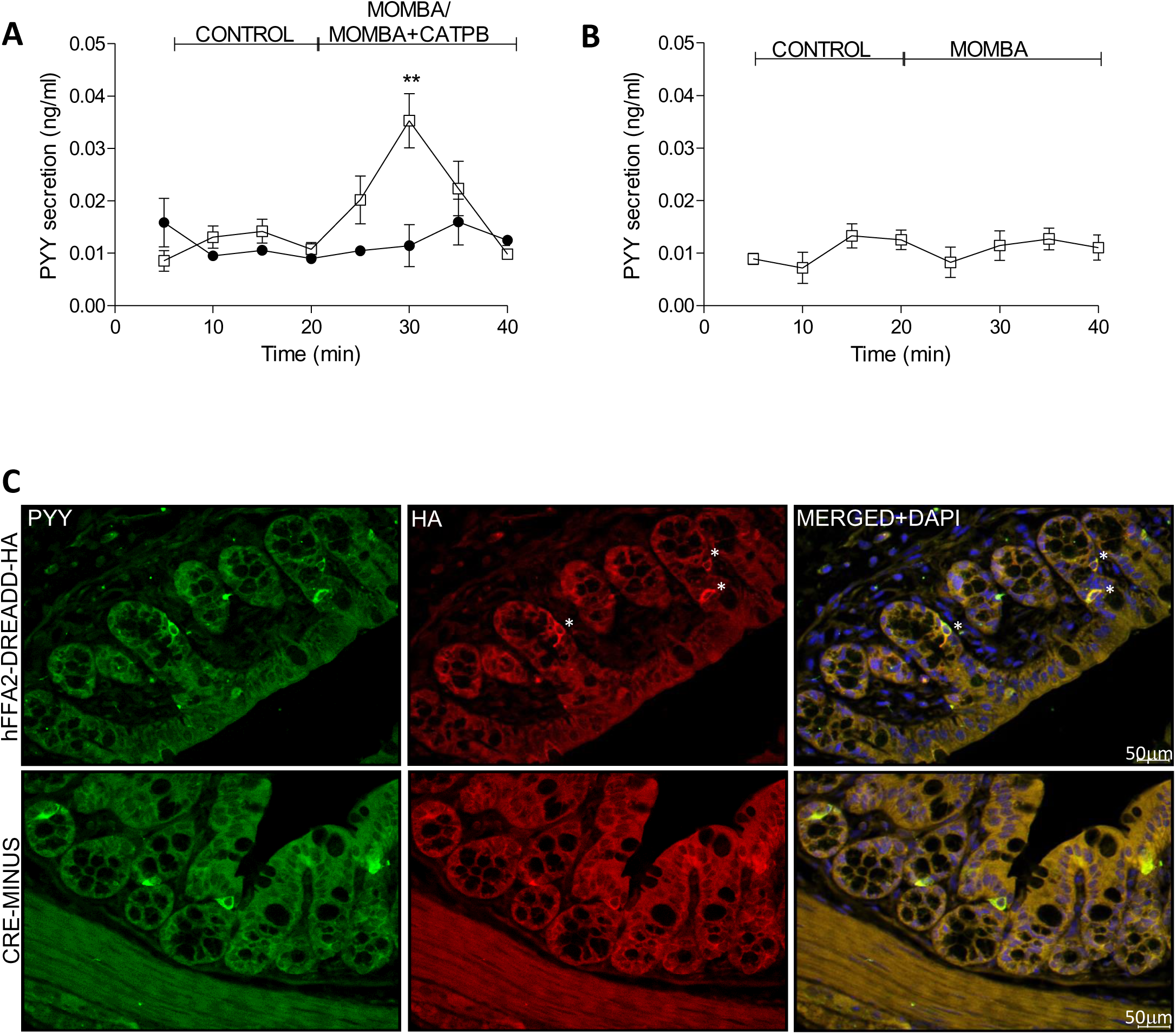
MOMBA-mediated activation of hFFA2-DREADD-HA promotes release of PYY. Following a period of flow of buffer into colonic tubes (**control**) introduction of MOMBA (**open symbols**) promoted transient PYY release from tissue of hFFA2-DREADD-HA expressing mice (** P < 0.01)(**left hand panel**) and this was prevented by co-exposure to CATPB (**filled symbols**) (**A**). MOMBA (**open symbols**) did not promote release of PYY from colonic tissue isolated from CRE-MINUS animals. Data represent means +/- S.E.M. from 4 (**A**) or 5 (**B**) different preparations. (**C**) Immunostaining of colonic sections from hFFA2-DREADD-HA (**top panels**) or wild type C57BL/6 (**bottom panels**) with either anti-PYY (**green, left**) or anti-HA (**red, middle**) identified subsets of PYY-expressing cells that were also positive for the hFFA2-DREADD-HA receptor (**stars**). In merged images (**right**) co-staining with DAPI (**blue**) identified cell nuclei.

### Electrophysiological recordings

We next wished to assess roles of SCFAs within the enteric nervous system. Afferent nerve fibres connect the colon to the spinal cord via dorsal root ganglia (DRGs). We therefore recorded action potentials from such afferents whilst perfusing the inside of either proximal colon or ileum (**Figure 4A**) from wild type mice. Perfusion of tissue with C3 resulted in a markedly increased (P < 0.05) rate of action potential generation in both preparations (**Figure 4B**).

**Figure 4.**
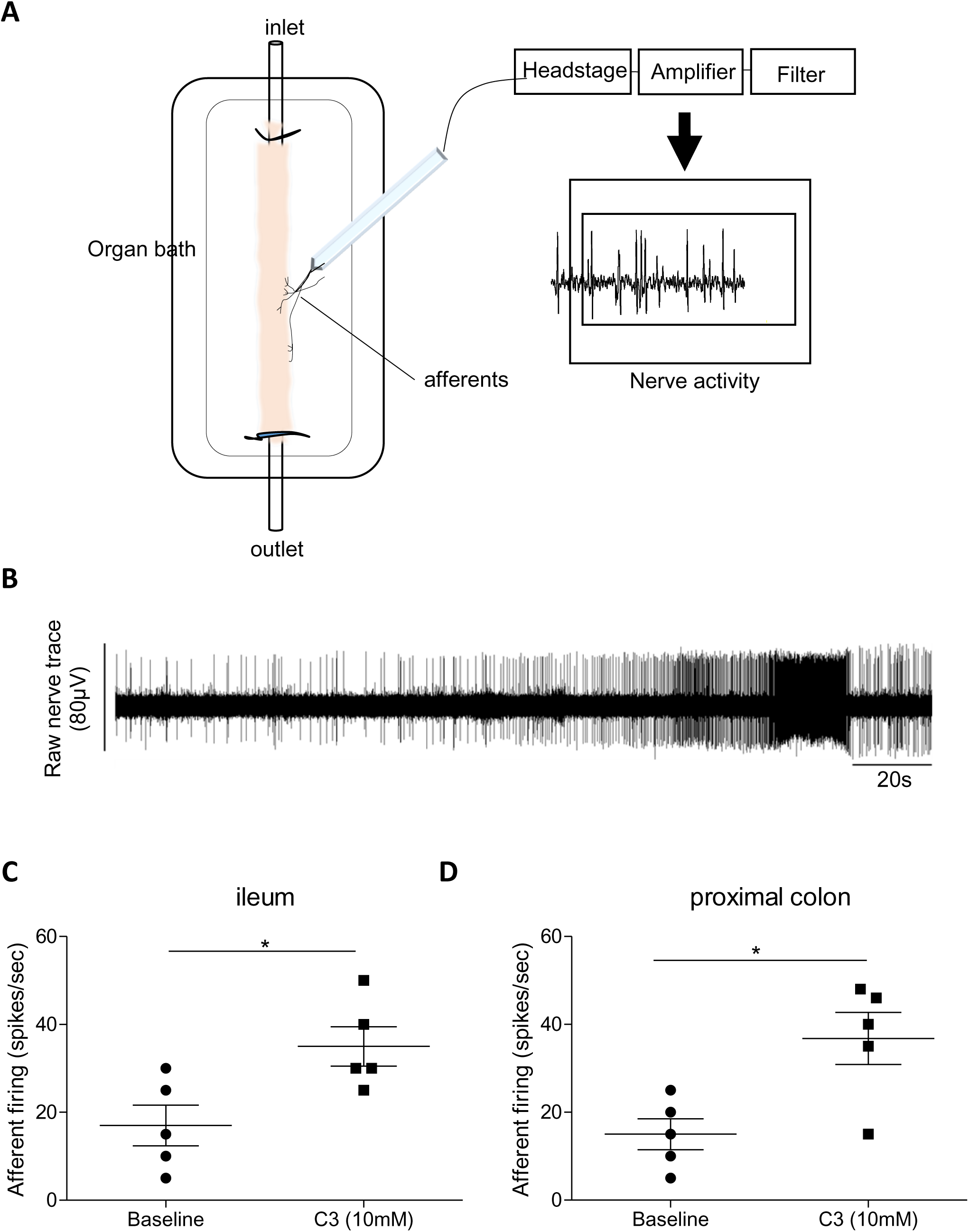
Colonic SCFAs promote firing of afferent nerves. A. Set up of recordings of electric activity of afferent nerve bundles. B. C3 (10 mM) was infused into intestinal segments taken from ileum or colon of wild type mice and maintained in an organ bath. Action potentials were measured over time from afferent nerve bundles associated with each region. The trace shown (**top**) is from a representative experiment conducted with tissue from colon. C3 increased afferent firing in both colon and ileum (* P < 0.05).

### Dorsal root ganglia express both FFA2 and FFA3

The expression of FFA3 in dorsal root ganglia (DRG) of mouse has previously been reported by Nohr et al., (29) using a FFA3-Red Fluorescent Protein (RFP) reporter mouse line in which expression of RFP, instead of the FFA3 receptor, is driven by *ffar3* gene promoter sequences. This is also the case for nodose vagal ganglia (29) and superior cervical ganglia in mice (35). Limited information on potential expression of FFA2 in such ganglia is available. Won et al., (36) reported significant expression of mRNA encoding FFA2 (and FFA3) in the celiac-superior mesenteric ganglion complex of rat but that FFA2 mRNA was not detectable in either nodose ganglia or DRGs. We initially examined this in colonic innervating DRGs that had been isolated from the T9-L2 region of the spinal cord of hFFA2-DREADD-HA expressing mice (**Figure 5**). Anti-HA staining was observed, and this was absent in equivalent preparations taken from CRE-MINUS mice (**Figure 5**). Whilst a limited extent of overlap of anti-HA immunostaining was observed with that for the neuronal marker PGP9.5, which was both widely distributed and detected in tissue from both hFFA2-DREADD-HA expressing and CRE-MINUS mice (**Figure 5**), the most intense anti-HA staining was in cells that interspersed between the PGP9.5-positive neurons (**Figure 5**).

**Figure 5.**
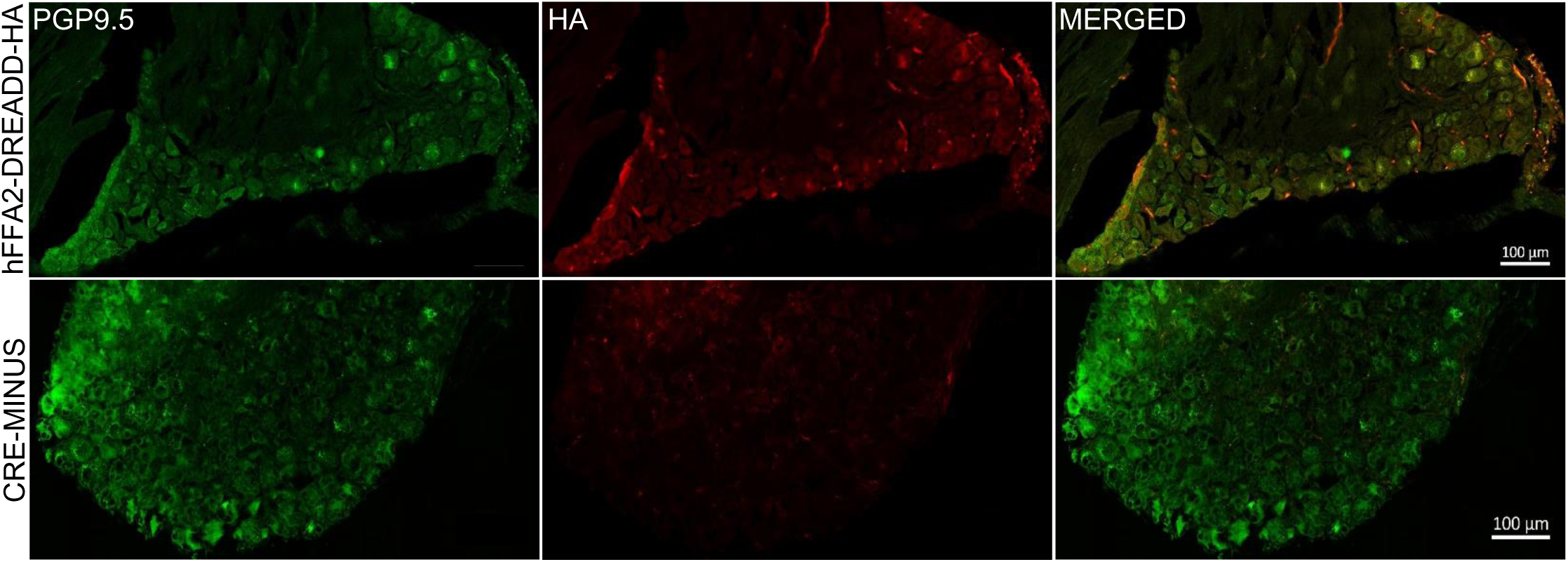
FFA2 is expressed in cells of dorsal root ganglia. Sections of dorsal root ganglia taken from either hFFA2-DREADD-HA expressing **(upper panels)** or CRE-MINUS **(lower panels)** were immunostained with anti-PGP9.5 to identify neurons (**left hand panels, green**) or with anti-HA to detect the receptor (**middle panels, red**). Merging of such images (**right hand panels**) showed modest co-expression but with additional anti-HA staining interspersed between neurons only in the hFFA2-DREADD-HA expressing sections.

### Elevation of Ca^2+^ in isolated dorsal root ganglion cells defines functional roles for both FFA2 and FFA3 receptors

To assess function of FFA2, cells from such DRGs were dispersed and plated onto matrigel-coated coverslips. After a recovery period cells were loaded with the Ca^2+^ indicator dye Fura-8-AM subsequent to mounting into an imaging chamber that allowed continuous perfusion of buffer. Addition of MOMBA (100 μM) to the buffer produced a rapid and partially sustained increase in intracellular [Ca^2+^] in a substantial proportion (41.4 +/- 10.0% mean +/- S.E.M., n = 103) (**Supplementary Table 1**) but not all, of the cells tested (**Figures 6A and 6B**). Washout of MOMBA and its replacement with C3 (1 mM) also resulted in elevation of intracellular [Ca^2+^] in most (82.1 +/-3.8%, mean +/- S.E.M., n = 84) (**Supplementary Table 2**) of the cells tested (**Figure 6A**). This was sometimes noted directly in the same cells that had responded previously to MOMBA (**Figure 6A**) but more broadly, however, in both a larger proportion of the cells and in a more robust manner than produced by MOMBA (see **Figure 6A** **and compare Figures 6B and 6C**). Following washout of either MOMBA or C3 the Ca^2+^ ionophore ionomycin was then applied to demonstrate integrity of the cell population. Here almost all cells tested produced a large increase in Ca^2+^ (**not shown**). Once more the effect of MOMBA was clearly an ‘on-target’ effect of the ligand at hFFA2-DREADD-HA because responses to MOMBA were essentially eliminated when cells were pre-incubated with the human FFA2 receptor specific antagonist CATPB (**Figures 6B** **and Supplementary Table 1**). By contrast, effects of C3 on cells within the population were not blocked by CATPB (**Figure 6C** **and Supplementary Table 2**). Moreover, the proportion of cells that responded to C3 was not different in DRGs isolated from CRE-MINUS animals (**Supplementary Figure 5**) whilst no effect of MOMBA was recorded in cells from CRE-MINUS mice (**Supplementary Figure 5**). Although the responses to C3 in DRG-derived cells from hFFA2-DREADD-HA expressing mice were entirely consistent with these effects being mediated via FFA3 we wished to produce further evidence to support this assertion and therefore employed the novel FFA3 allosteric agonist TUG-1907 (22). This ligand produced substantial elevation of Ca^2+^ in isolated DRG-derived cells (**Figure 6D**). As this ligand did not produce such effects in DRGs isolated from FFA3 knock-out mice (**Figure 6D**), this confirmed the presence and functionality of FFA3.

**Figure 6.**
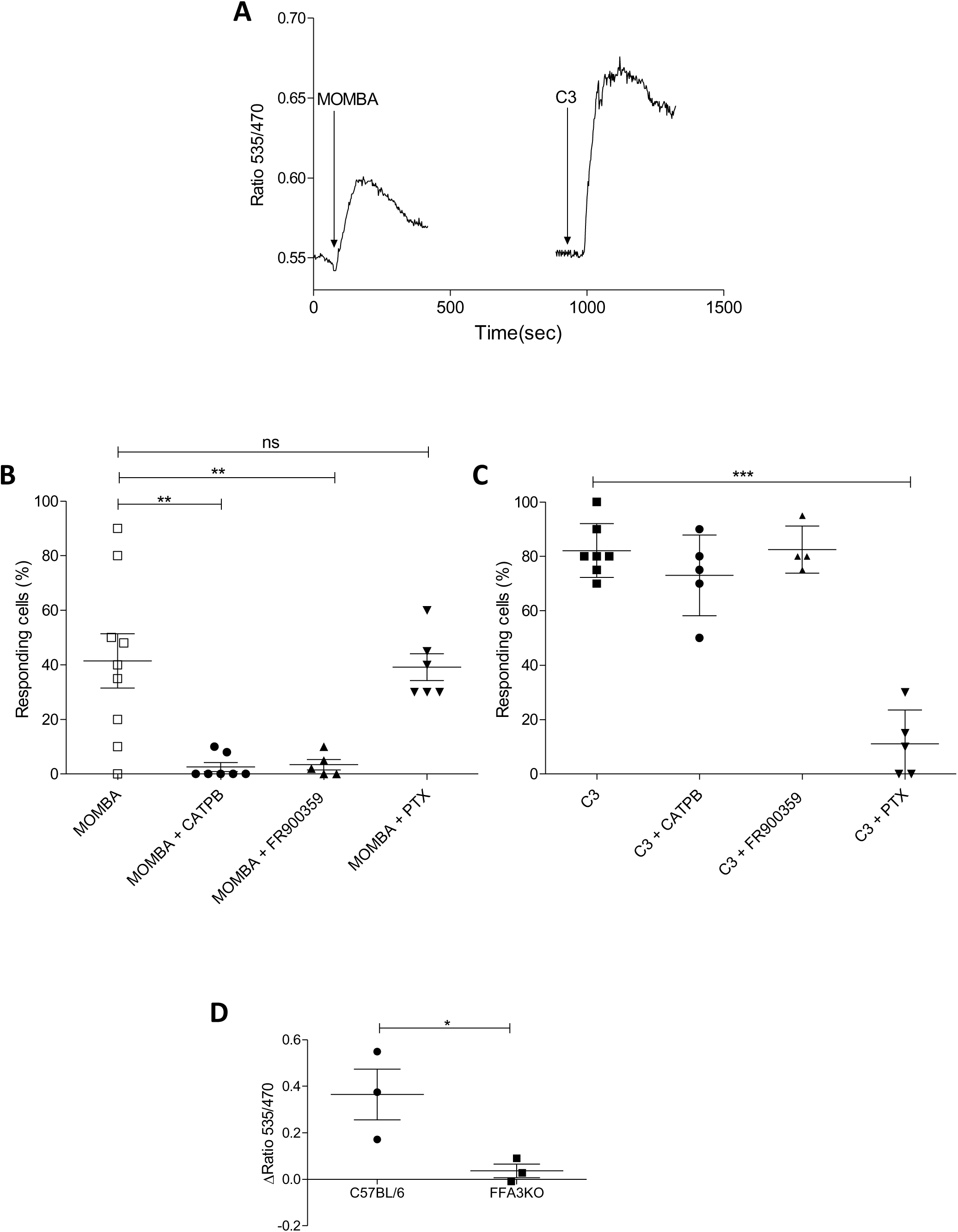
Both FFA2-DREADD and FFA3 are functional in cells of dorsal root ganglia but elevate Ca^2+^ by different mechanisms. Single cell Ca**^2+^** imaging studies were performed on cells isolated from DRGs taken from hFFA2-DREADD-HA expressing mice. **A**. Individual cells were exposed sequentially to MOMBA and then C3 with washout of MOMBA prior to treatment with C3. A representative example is shown. **B**. Cells taken from mice were exposed to MOMBA or, from 7 mice to MOMBA + CATPB. CATPB blocked the effect of MOMBA (** P < 0.01). Cells were also exposed to MOMBA after pre-treatment with the G_q_/G_11_ inhibitor FR900359 (for 15 minutes) or pertussis toxin (for 24 hours). FR900359 blocked the effect of MOMBA (** P < 0.01) but pertussis toxin treatment did not. **C**. Studies akin to **B** were conducted with C3 as a potential FFA3 activator. The effect of C3 was blocked by pre-treatment with pertussis toxin (*** P < 0.001) but unaffected by either CATPB or FR900359. Data represent the mean % of cells prepared from tissue of individual mice that responded to ligand treatments. **B** and **C** One-way analysis of variance followed by Bonferroni’s Multiple Comparison Test. **D**. The selective FFA3 activator TUG-1907 (5 μM) promoted Ca^2+^ elevation in cells from wild type (WT) but not (* P < 0.05, unpaired t-test) in cells taken from FFA3 knock-out (FFA3KO) animals. Peak increases in Ca^2+^ are shown.

### Mechanisms of FFA2- and FFA3-mediated Ca^2+^ elevation in cells of dorsal root ganglia

The ability of MOMBA to promote elevation of Ca^2+^ levels in DRG-derived cells taken from hFFA2-DREADD-HA mice was blocked by co-incubation with the Gα_q_/Gα_11_ inhibitor FR900359 (37) (**Figure 6B** **and Supplementary Table 1**). This is consistent with the cation being released from inositol 1,4,5 trisphosphate (IP_3_)-sensitive internal stores. By contrast effects of MOMBA were unaffected by pre-treatment with pertussis toxin (**Figure 6B** **and Supplementary Table 1**) which causes ADP-ribosylation of Gα_i_ family subunits. Very differently, responses to C3 were instead blocked by pre-treatment with pertussis toxin, but not by treatment with FR900359 (**Figures 6C** **and Supplementary Table 2**). This is consistent with the effect of C3 being mediated by FFA3 in these cells and proceeding via release of β/γ complexes from G_i_α-containing G protein heterotrimers (38–39). These results indicate that activation of both FFA2 and FFA3 results in elevation of Ca^2+^ in DRG-derived cells, but that the two receptors engage with different G proteins and use distinct mechanisms to mediate these effects.

### Nodose ganglia also express FFA2

Anti-HA staining of nodose ganglia isolated from hFFA2-DREADD-HA mice showed a pattern similar those observed in DRGs with clear staining that overlapped only to a limited degree with the neuronal marker PGP9.5 (**Figure 7A**). Moreover, as in cells isolated from DRGs, single cell Ca^2+^ imaging of isolated cells from nodose ganglia from these animals showed that only a proportion of them (37.2 +/- 8.4%) responded to addition of MOMBA (**Figure 7B**) whereas a markedly larger proportion (86.9 +/- 6.5%) responded to C3 (**Figure 7B**). Again, like in DRG-derived cells no response to MOMBA was observed when cells were exposed to CATPB in addition to MOMBA (**Figure 7B**). Once more this confirms that the effect of MOMBA does indeed reflect activation of hFFA2-DREADD-HA and, therefore, that this subset of cells derived from nodose ganglia express functional hFFA2-DREADD-HA. By contrast, in equivalent cells isolated from wild type mice no response to MOMBA was recorded (1.6 +/- 1.0%, 59 cells, n = 5 animals) (**Figure 7C**), whilst 85.0 +/- 5.5% of the cells (51 cells, n = 5 animals) responded to C3 (**Figure 7C**) and this was not different from the proportion of cells able to respond to C3 isolated from hFFA2-DREADD-HA expressing mice (**Figure 7B**).

**Figure 7.**
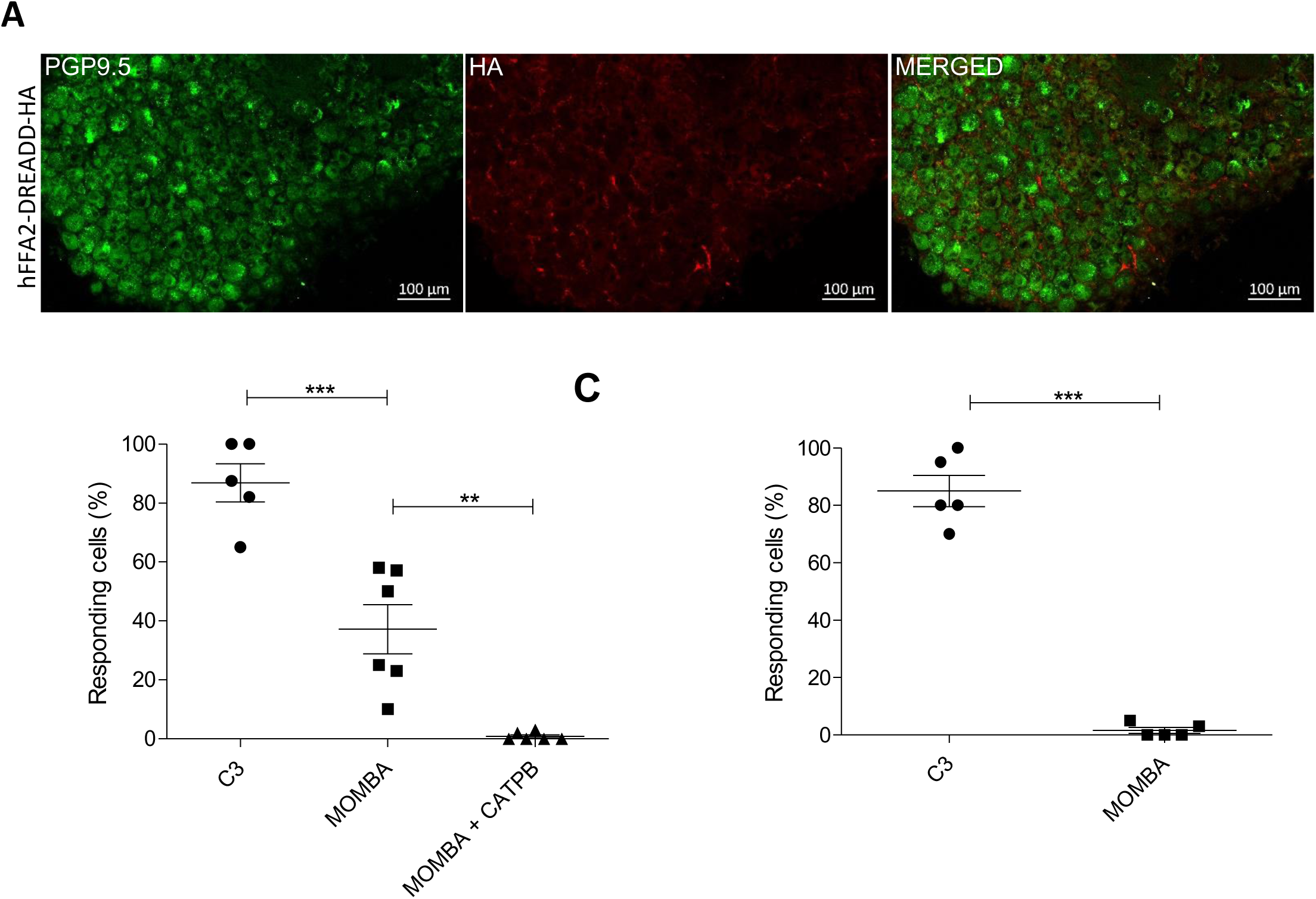
FFA2 is also expressed and functional in nodose ganglia of mice. Sections of nodose ganglia from FFA2-DREADD-HA expressing mice were immunostained with anti-PGP9.5 to identify neurons (**left hand panel, green**) or with anti-HA to detect the receptor (**middle panels, red**) (**A**). Merging of such images (**right hand panel**) showed modest co-expression, but with additional anti-HA staining interspersed between neurons in the hFFA2-DREADD-HA expressing sections. **B**. Dissociation and plating of cells from nodose ganglia from either hFFA2-DREADD-HA (**B**) or wild type C57BL/6 (**C**) mice was followed by single cell Ca^2+^ imaging studies as in Figure 6. As in cells dissociated from DRG C3 generated responses in equal proportions of cells of the two lines, MOMBA was active only in cells produced from hFFA2-DREADD-HA expressing mice, and this effect of MOMBA was blocked by co-treatment with CATPB (** P <0.01). A greater fraction of cells from hFFA2-DREADD-HA expressing animals responded to C3 than to MOMBA (*** P < 0.001). Data represent the mean % of cells prepared from tissue of individual mice that responded to ligand treatments.

### Gut short chain fatty acid receptors promote activation of neurons in the dorsal horn of the spinal cord

Finally, we wished to assess whether short chain fatty acids present in the gut might promote central nervous system stimulation and, if so, if this was a GPCR-mediated effect. To do so, initially we perfused either saline or C3 via the rectum into the colon of anaesthetized wild type mice. Following sacrifice 2 hours later activation of neurons in the dorsal horn of the spinal cord was assessed by measuring induction of expression of the early immediate gene c-fos (**Figure 8A, compare Figures 8B and 8C**). C3 produced a substantial increase in c-fos positive neurons compared to saline (P < 0.05) (**Figure 8A**). To dissect the contribution of FFA2 to this effect we performed equivalent experiments in hFFA2-DREADD-HA expressing mice. Here, introduction of MOMBA produced a marked increase (P < 0.01) compared to saline infusion in the number of c-fos positive neurons (**Figure 8A, compare Figures 8D and 8E**). By contrast MOMBA did not promote such an effect in the CRE-MINUS animals (**Figures 8A and 8F**), confirming this effect of MOMBA to reflect activation of the hFFA2-DREADD-HA receptor. Interestingly C3 **also/did not** promote stimulation of c-fos expression in the hFFA2-DREADD-HA expressing animals, suggesting a potential additional role for FFA3. Experiments with TUG-1907 also **resulted/did not result** in an increase in neurons positive for c-fos expression, providing further evidence that activation of FFA3 at the level of the gut/enteric nervous system initiates stimulation within the spinal cord (**Figure 8**).

**Figure 8.**
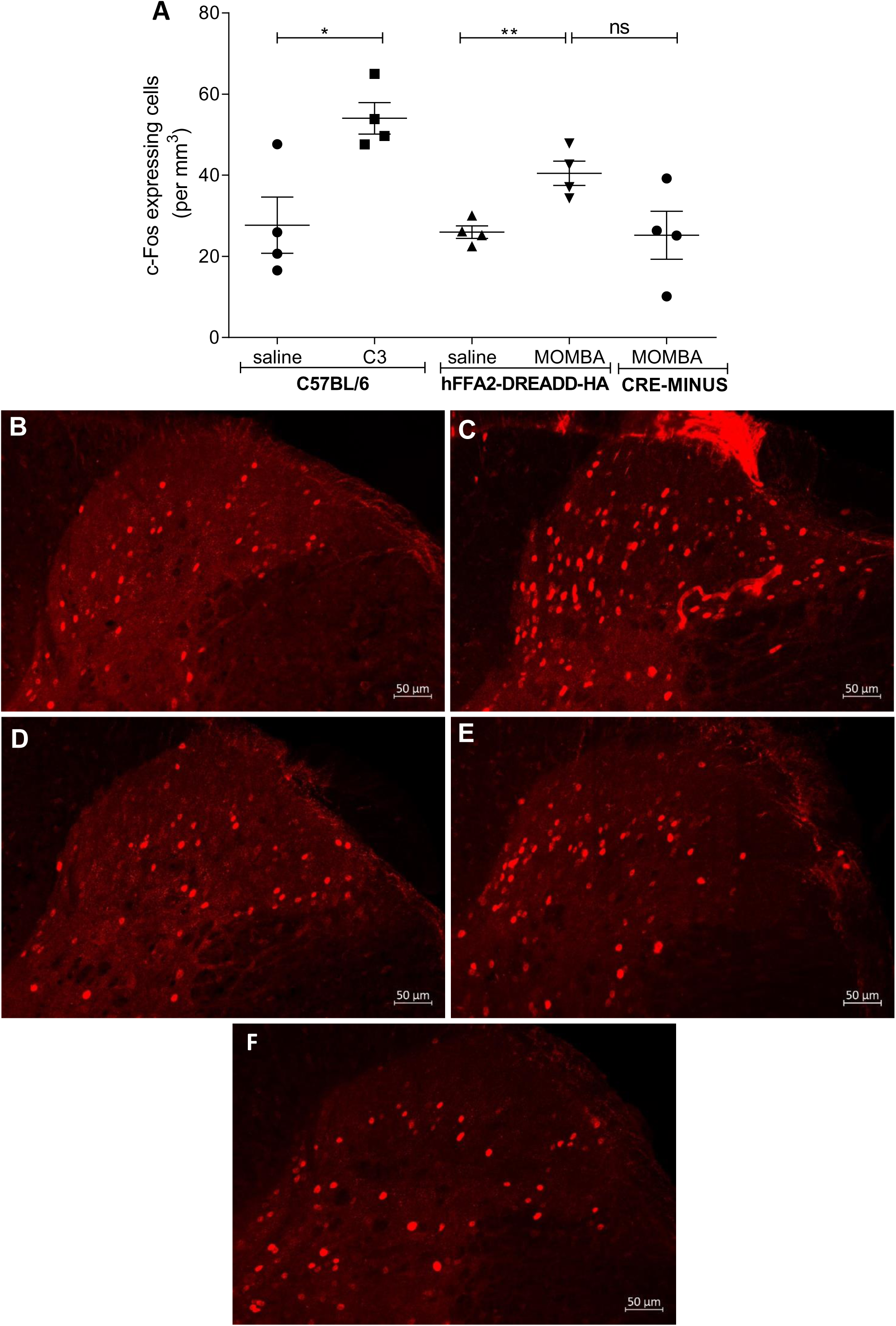
Gut short chain fatty acid receptors promote activation of neurons in the dorsal horn of the spinal cord. 2 hours after introduction of saline, C3 or MOMBA into the colon of wild type C57BL/6, hFFA2-DREADD-HA or CRE-MINUS mice the number of c-fos expressing neurons in the dorsal horn of the spinal cord was quantified (**A**). C3 increased c-fos expressing cells in wild type (* P < 0.05), MOMBA also did so in hFFA2-DREADD-HA expressing (** P < 0.01) but not CRE-MINUS mice. Unpaired t-test. See also Supplementary Table 3. Representative images of C57BL/6 plus saline (**B**), or plus C3 (**C**); hFFA2-DREADD-HA plus saline (**D**), or plus MOMBA (**E**); CRE-MINUS plus MOMBA (**F**) treatments are shown for illustration.

## Discussion

The introduction of DREADDs, derived from members of the muscarinic acetylcholine family of GPCRs, into both isolated cell types and intact animals has provided multiple new insights into functions regulated by signalling cascades controlled by these receptors when the DREADD selective ligand CNO is provided (40–42). This has become a widely used approach in many areas of, particularly, neurobiology (43). Much less attention has been given to applying the DREADD concept to help define specific functions of a GPCR at which the endogenous ligand(s) for the wild type receptor also activates other, closely related and sometimes co-expressed GPCRs. Previously we demonstrated that by introduction of two amino acid mutations the human FFA2 receptor lost binding and responsiveness to SCFAs, but in parallel gained responsiveness to sorbate (18). Because it has remained unclear in both *ex vivo* and *in vivo* studies if various functions of SCFAs are mediated by FFA2, or are potentially mediated by the related receptor FFA3 or, indeed, by other GPCRs that can be activated by SCFAs (44), we recently developed a transgenic knock-in line of mice in which mouse FFA2 was replaced by the above described humanized FFA2-DREADD (19). Because of the well appreciated lack of specificity of many anti-GPCR antisera we also added the HA-epitope tag sequence to the C-terminus of the hFFA2-DREADD construct to allow immunolocalization of the expressed receptor in these animals (19). Importantly, as well as demonstrating expression of hFFA2-DREADD-HA in cells and tissues where expression of mFFA2 was anticipated, herein this has allowed us to clearly demonstrate expression of the receptor in a range of cells and tissues within the gut-brain axis, including cells that express and can release the enteroendocrine hormone PYY and cells of both the nodose and dorsal root ganglia. The presence of FFA2 within such ganglia has been unclear (36) but now as well as the receptor being detected we have also shown the functionality of FFA2 in subsets of cells isolated from these ganglia, based on their responsiveness to the novel hFFA2-DREADD specific agonist identified in these studies, MOMBA. Moreover, because such effects of MOMBA were blocked by co-application of the hFFA2-DREADD active antagonist CATPB this confirms the effects of MOMBA to be ‘on target’. These studies provide the first experimental evidence of the capacity of FFA2 to generate signals in cells of these ganglia. Of key importance the strategy for ‘knock-in’ transgenic expression of hFFA2-DREADD-HA was developed to ensure uninterrupted expression of other well defined SCFA receptor FFA3 (19), the gene for which is closely linked with that for FFA2 on chromosome 7 in mouse (9). Although this receptor has previously been shown to be expressed in such ganglia (29–30, 35), because the hFFA2-DREADD-HA receptor is not able to bind or be activated by SCFAs, then signalling cascades initiated by C3 in cells and tissue of these animals can be predicted to be mediated by FFA3. Previously, selective activation of FFA3 has been difficult to define pharmacologically because of the activity of SCFAs also at wild type FFA2. However, to more fully define effects generated by FFA3 we have also used a novel, highly FFA3 selective agonist, TUG-1907, that we have recent identified and characterized (22).

In our previous studies using the hFFA2-DREADD-HA expressing mice we employed sorbate as selective agonist of this receptor (19). It is clearly beneficial for studies at any potential therapeutic target to have more than a single agonist ligand. Although sorbate has extremely low toxicity, it is widely used as a food preservative (21). Because of its potential anti-bacterial and anti-fungal effects which might potentially affect longer-term studies in which we provide sorbate in the drinking water of these animals to identify other, chemically distinct, hFFA2-DREADD agonist ligands we initially screened a selected library of 1210 small molecule compounds with relatedness to sorbate. This identified MOMBA as having appropriate characteristics as a selective agonist. Moreover, as both 4-methoxy-3-chloro-benzoic acid and 4-methoxy-3-hydroxy-benzoic acid were subsequently also identified from a further, more targeted, compound set as actives at hFFA2-DREADD, MOMBA was taken into broader ranging studies. Importantly, MOMBA was shown to be as efficacious as sorbate and somewhat more potent across a range of assays. Moreover, like sorbate MOMBA was confirmed to be an orthosteric agonist at hFFA2-DREADD and therefore binds within the same pocket as SCFAs do at wild type forms of FFA2 (9, 12). This implied that effects of MOMBA acting at the receptor should be fully blocked, and in a competitive manner, by the human FFA2 receptor selective antagonists CATPB and GLPG0974. This was the case, and such in cells and tissues from the hFFA2-DREADD-HA expressing animals blockade of effects of MOMBA by either of these antagonists provides further certainty that effects of MOMBA are indeed mediated by the receptor.

Key outcomes of the studies reported herein were that in both nodose and dorsal root ganglia both FFA2 and FFA3 are expressed and that both are functional. However, although selective activation of either FFA2 or FFA3 resulted in elevation of Ca^2+^ it is remarkable that they did so by completely different mechanisms. It is well appreciated that FFA3 couples highly selectively to members of the pertussis toxin-sensitive G_i_-family of G proteins (24–25). As such, it was anticipated that in cells and tissues from hFFA2-DREADD-HA expressing mice FFA3-mediated effects should be blocked by pre-treatment with pertussis toxin. An established mechanism to allow G_i_-coupled receptors to elevate Ca^2+^ is via β/γ complexes released by activation of G_i_α-containing heterotrimers (38–39). By contrast FFA2 (24–25), and indeed the hFFA2-DREADD construct (19), is able to promote activation and downstream signalling of a wide range of G proteins, at least in transfected cell lines. Moreover, we have previously shown that in tissues of the hFFA2-DREADD-HA expressing mice, regulation of lipolysis is transduced by G_i_-family G proteins whilst release of the hormone GLP-1 requires activation of G_q_/G_11_ family G proteins (19). Herein, Ca^2+^ elevation in cells of DRGs by FFA2 was transduced entirely by G_q_/G_11_ as agonist effects were abolished by pre-treatment with the selective G_q_/G_11_ inhibitor FR900359 (37) but completely unaffected by pre-treatment with pertussis toxin. How the hFFA2-DREADD-HA receptor selects between co-expressed G proteins remain unclear but will be a topic for future analysis.

Perhaps most remarkable amongst the outcomes reported herein is the capacity for short chain fatty acid receptors that are activated by agonists introduced into the colon to cause activation of not only afferent nerve bundles in the enteric nervous system but to promote activation of neurons at the level of the dorsal horn of the spinal cord. This was recorded by measuring turn-on of expression of the early immediate gene c-fos (45–46) in such neurons within 2 hours of provision of ligands into the colon of anaesthetized mice. These studies provide clear evidence of the role of gut microbiota-derived SCFAs to co-ordinate and control responses between the gut and the central nervous system. Given the rapidly emerging evidence that the microbiota and their dysbiosis can be key contributors to a variety of long-term central nervous system deficits and malfunctions (47–50) the current studies provide insights into direct chemical links via the gut brain axis produced by SCFAs that are the most abundant products of fermentation of fibre, and more generally of metabolism, by the microbiota. Whilst circulating SCFAs can cross the blood-brain barrier directly and may influence the activity of microglia involved in neuroinflammation via such a systemic route (48) it is unclear if concentrations of SCFAs in the circulation would routinely be sufficiently high to achieve substantial activation of receptors such as FFA2 and FFA3 at that level. By contrast, as well as providing entirely new insights into the roles of FFA2 and FFA3, these studies provide fascinating hints as to how the high concentrations of SCFAs generated in the gut may transmit such signals to the brain via a neuronal gut-brain relay. It therefore may be possible to target the gut-brain axis in the therapeutic context of various CNS disorders by targeting FFA2 and/or FFA3 and we provide mouse models to further dissect and assess these processes and opportunities.

## Materials and Methods

### Compounds

Compounds for screening were selected in two stages from the AstraZeneca compound collection. First, a larger set of 1210 carboxylic acids was assembled to generate an initial list of actives by screening. The first set of compounds aimed to balance (i) similarity to known agonists (including sorbate), (ii) chemical diversity, and (iii) physicochemical properties consistent with favourable ADME (absorption, distribution, metabolism, and excretion) profiles of molecules. To this end, carboxylic acids were identified by substructure matching, and compounds with either (i) molecular weight > 350 Da, (ii) calculated octanol-water partition coefficient logP > 3.5, (iii) calculated polar surface area > 110 Å^2^, or (iv) unwanted chemical groups were removed. After clustering, a diverse subset of 1210 compounds was manually selected. The second stage of compound selection involved identifying close analogs to the active molecules in the first screen. Thus, 320 compounds were selected using multiple substructure searches. Subsequent batches of MOMBA were purchased from FLUOROCHEM, Hadfield, United Kingdom.

### β-arrestin-2 recruitment assays

β-arrestin-2 recruitment to a receptor of interest was assessed using a bioluminescence resonance energy transfer (BRET)-based assay. HEK-293T cells were co-transfected with the desired receptor tagged with eYFP at its C terminus, and with β-arrestin-2-*Renilla* luciferase (ratio 4:1), using 1 mg.ml^-1^ PEI, linear MW-25000 (ratio 1:6 DNA/PEI). Subsequent experimental procedures were performed as described previously (19). The final concentration of coelenterazine-h used as substrate was 5 µM. BRET measurements were performed using a PHERAstar FS reader (BMG-Labtech, Offenburg, Germany). The BRET ratio was calculated as emission at 530 nm/emission at 485 nm. Net BRET was defined as the 530 nm/485 nm ratio of cells co-expressing *Renilla* luciferase and eYFP minus the BRET ratio of cells expressing only the *Renilla* luciferase construct in the same experiment. This value was multiplied by 1000 to obtain mBRET units.

### Primary compound screening

Compounds were assayed in a single point BRET assay at 100 μM containing 1 % DMSO. Basal wells containing assay buffer with 1% DMSO, and positive stimulation wells containing 1 mM sorbate were included in all plates. Each plate contained two ‘*Renilla* luciferase’ points containing cells lacking receptor expression. Data from primary screening were analysed using Microsoft Excel software and activities of the compounds were calculated using the following formula: Activity (%) = (mBRET compound – mBRET basal) / (mBRET stim – mBRET basal) x100, where mBRET compound is the mBRET value obtained from wells treated with the test compound, mBRET basal is the average of the mBRET values obtained from wells treated with assay buffer and mBRET stim is the average of the mBRET values obtained from cells treated with 1 mM sorbate. Compounds were considered to be possible hits if the activity was higher than the mean + 3×S.D of the overall activity in the whole assay. Hits were considered to be confirmed if the activity remained over this threshold in a second independent assay. Reliability of the assay was estimated by calculating Z′ values (ref 23) for each plate, using the formula: Z’ = 1 - {[3 x σstim) + (3 x σbasal)]/(µstim - µbasal)} where σstim and σbasal are the S.D. values of wells containing 1 mM sorbic acid and assay buffer respectively and μstim and μbasal are the means for wells containing 1 mM sorbate and assay buffer respectively.

### Cell lines

HEK-293T cells were maintained in Dulbecco’s Modified Eagle’s Medium (DMEM) supplemented with 0.292 g.l^-1^ L-glutamine, penicillin/streptomycin mixture and 10% (v/v) fetal bovin serum (FBS) at 37 °C in a 5% CO_2_ humidified atmosphere. For experiments using transiently transfected HEK-293T cells, transfections were carried out using 1 mg.ml^-1^ polyethyleneimine (PEI) (MW-25000) and experiments conducted 48 h post transfection.

Flp-In™ T-REx™-293 cells (Invitrogen) were maintained in DMEM without sodium pyruvate, supplemented with 10% (v/v) FBS, 1% penicillin/streptomycin mixture, and 10 µg.ml^-1^ blasticidin at 37 °C in a 5% CO_2_ humidified atmosphere. To generate Flp-In T-REx 293 cells able to inducibly express the various FFA receptor constructs, cells were co-transfected with a mixture containing the desired cDNA in pcDNA5/FRT/TO vector and pOG44 vector (1:9) using PEI. Transfected cells were selected using 200 μg.ml^-1^ hygromycin B. Expression of the appropriate construct from the Flp-In™ T-REx™ locus was induced by treatment with 100 ng.ml^-1^ doxycycline for 24 h.

### Cell signalling assays

Each of regulation of forskolin-amplified levels of cAMP, production of inositol monophosphates (IP_1_) and binding of [**^35^**S]GTPγS was performed as detailed in (19).

### Animal maintenance

The generation and characterization of both transgenic FFA2-DREADD-HA expressing and CRE-MINUS mouse lines is detailed in (19). All animals were bred as homozygous onto a C57BL/6N background. Male and female mice were used in this study unless otherwise stated. Mice were fed *ad libitum* with a standard mouse chow diet. Maintenance and killing of mice followed principles of good laboratory practice in accordance with UK national laws and regulations. All experiments were conducted under a home office licence held by the authors.

### Immunohistochemistry

Colonic tissues were isolated, mounted and processed using previously described (19) methods. Sections were immunostained with anti-HA (1:100; Sigma-Aldrich anti-HA high affinity-clone 3F10) and anti-PYY (1:400 Abcam-Ab1 ab22663) antibodies and mounted with VECTASHIELD® Vibrance™ Antifade Mounting Medium with DAPI (Vector Laboratories). Images were taken with an EVOS M7000 Imaging System (ThermoFisher Scientific).

### DRG and nodose ganglia isolation and calcium imaging

Colonic innervating DRGs were isolated from the T9-L2 region of the spinal cord of wild type and transgenic animals and immediately placed in cold Hanks′ balanced salt solution (HBSS; Sigma-Aldrich). Isolated DRGs were initially digested with HBSS containing L-cysteine (0.3 mg.ml^-1^) and papain (2.0 mg.ml^-1^) for 20 min at 37°C. The solution was removed and replaced with HBSS contain collagenase (4.0 mg.ml^-1^) and dispase (4.0 mg.ml^-1^) (20 min at 37°C) for further digestion. The collagenase solution was then replaced with DMEM to stop the reaction. The DRGs were finally dissociated by mechanical trituration using a pipette. Dissociated cells were plated on matrigel-coated coverslips and placed in an incubator (37°C and 5% CO_2_). Following a 2 h incubation cells were flooded with 90% DMEM (Sigma) supplemented with 10 % fetal calf serum and 1 % PenStrep and further incubated overnight at 37°C and 5 % CO_2_.

To measure intracellular calcium and its potential regulation, dissociated cells on the coverslips were loaded with Fura 8-AM (3 µM) (Stratech Scientific Limited) for 20 min at 37°C in the dark. Coverslips were then placed in a recording chamber and mounted onto a invert fluorescent microscope (Nikon TE2000-E; Nikon Instruments, Melville, NY) equipped with a (NA = 1.3) oil-immersion Super Fluor objective lens (x40), an Optoscan monochromator (Cairn Research, Faversham, Kent, UK) and a digital Cool Snap-HQ CCD camera (Roper Scientific/Photometrics, Tucson, AZ). Illumination of the preparation was achieved by a Meta Fluor imaging software (Molecular Devices, San Jose CA, version 7.8.8). Clusters of cells were randomly selected for real time imaging and continuously perfused with HEPES buffer (composition: HEPES 10 mM, NaCl 135 mM, glucose 10 mM, KCl 5 mM, CaCl_2_ 2 mM and MgCl_2_ 1 mM, pH 7.4) for 20 min at room temperature. All test ligands were diluted in HEPES buffer and perfused through the chamber for 3 min, followed by a final application of the Ca^2+^ ionophore ionomycin (5µM), as a positive control. Results are expressed as relative fluorescence (RF), *n* numbers are presented as N = number of mice and n = number of cells. Similar methods were used for cell from nodose ganglia.

### Detection of c-fos expression in spinal cord

T9–T10 sections (40 µM) were cut on a vibrating blade microtome Leica VT1200 or VT1000S, Leica), with peroxidase quenching buffer (2% H_2_O_2_ in PBS) for 30min (RT), followed by 50% ethanol (30min, RT). Sections were subsequently incubated with primary antibody c-fos (goat 1:500, Santa Cruz) for 72h, followed by overnight incubation with biotinylated secondary antibody (1:500, Jackson Laboratory). c-fos staining was detected using TSA tetramethylrhodamine Kit (NEL702001KT, Perkin Elmer). Sections were washed in PBS (3×5min) and then incubated with Strepavidin for 4h (1:100, RT). The sections were again rinsed with PBS (3×5min), followed by an incubation period with tyramide solution [1:50] in amplification buffer (0.0015% H_2_O_2_) for 7min (RT). The tissues were then rinsed with PBS (3×5min). Sections were mounted on glass slides in Vectashield anti-fade mounting medium with DAPI.

### Colon stimulation

Vehicle or test compounds were administered via enemas (200 μl). Colorectal stimulation was performed by gently inserting a tube (2.5cm) into the rectum. The solution was slowly injected. Mice were allowed to recover and monitored for discomfort/pain for 2h, followed by transcardial perfusion fixation with 20 ml of PBS then 20 ml of 4% formaldehyde in a 0.1M phosphate buffer.

### Cryo-sectioning and immunostaining

Following cervical dislocation, DRGs were quickly dissected out and fixed in 4% PFA for 90 min. The specimens were cryo-sectioned at 30 µm and thaw-mounted onto adhesive slides (Leica). Slides were washed in Tris-buffered saline (TBS) containing 0.3% Triton X-100 and incubated with blocking buffer (TBS, 0.3% Triton X-100, 3% goat serum, 5% BSA) for 2h, followed by incubation with primary antibodies (rat-HA, 1:100, rabbit-PGP9.5, 1:1000) for 24h (4°C). Slides were further incubated for 2h (RTP) with secondary antibody 1:400 (Alexa 448 fluor goat anti-rat, or Alexa fluor 647 goat anti-rabbit). Slides were mounted with Vectashield mounting solution. Fluorescent images were visualized and captured with a Zeiss confocal microscope.

### In vivo gastrointestinal transit

GI transit was measured as previously described by (19). Briefly, a cohort of male (12-18 weeks) hFFA2-DREADD-HA, wild type and CRE-MINUS mice were single caged with free access to food and water. After one week of acclimatisation, mice were gavaged with a solution of carmine red (300 µl; 6 %; Sigma-Aldrich) suspended in methylcellulose (0.5 %; Sigma-Aldrich). Total GI transit time was measured as the time between oral gavage (Time=0) and the appearance of the first red pellet. Following the initial transit studies, half the mice from each group were randomly selected and provided with MOMBA (15 mM) in the drinking water, whereas the other half (control) continued drinking water without MOMBA. After one week the GI transit of all mice was again measured. In certain studies, MOMBA was then removed and the mice were again provided with water followed by a further gavage with carmine red a week later.

### Colonic crypt isolation

As previously described (19) the colon was immediately removed and placed in ice-cold Krebs solution. The colon was cut longitudinally and pinned on a sylgard coated dish. The muscle was gently removed and the remaining tissue was chopped using a scalpel. The tissue was then washed three times with cold PBS. For tissue digestion, the colon was placed in medium containing 0.3 mg.ml^-1^ collagenase XI (Sigma-Aldrich) for 15 min at 37°C. The supernatant was then collected and the remaining tissue was further digested with collagenase. This process was repeated two more times to allow complete digestion of the colon. Isolated crypts were plated on matrigel (Corning)-coated wells and incubated overnight at 37°C and 5 % CO_2_ in DMEM (25 mM glucose) supplemented with 10% fetal bovine serum, 1% glutamine and 1% Penicllin/Streptomycin. On the following day, all wells were washed with 138 buffer and challenged with test compounds. After two hours, the supernatants and lysates were centrifuged (4°C, 18,000 x g). Active GLP-1 secretion was measured by ELISA (Millipore).

### Organ bath studies

GLP-1 and PYY secretion from intact colon was investigated in hFFA2-DREADD-HA, CRE-MINUS and wild-type mice as previously described in detail (19). The entire colon was removed and placed in an purpose built organ bath (3 ml) perfused with carbogenated (95% O_2_–5% CO_2_) Krebs solution (composition, mM: NaCl 120; KCl 5.9; NaH_2_PO_4_ 1.2; MgSO4 1.2; NaHCO_3_ 15.4; CaCl_2_ 2.5; glucose 11.5) (34°C). The colon was attached from either end to an inlet and outlet port, allowing intraluminal perfusion (10 ml.hr^-1^) of vehicle or test ligands using a syringe pump (Sigma-Aldrich).

GLP-1 and PYY secretion was assessed by perfusing either Krebs or MOMBA (0.1 mM) through the lumen. To further assess the specificity of MOMBA, MOMBA was also applied in the presence of the human FFA2 specific antagonist CATPB (10 μM). In this case, CATPB was also applied 15 minutes before the co-application of MOMBA (0.1 mM) and CATPB (10 μM). During intraluminal perfusion, supernatants were collected every 5min. Total PYY (Phoenix) and active GLP-1 (Millipore) concentration was measured by ELISA.

### Data analysis and curve fitting

All data in this manuscript represent the mean ± S.E.M. or S.D. where noted of at least three independent experiments. Data analysis and curve fitting were conducted using the GraphPad Prism software package 8.1.0 (GraphPad). Concentration-response data were fit to three-parameter sigmoidal concentration-response curves. Statistical analyses were performed using a two-tailed t-test, one- or two-way ANOVA analyses followed by Dunnett or Bonferroni post hoc test as indicated. Allosteric parameters were calculated by using the operational model equation as described previously (19).

## Acknowledgements

The studies reported were funded by the Biotechnology and Biosciences Research Council (grant numbers BB/L027887/1 (to GM), BB/L02781X/1 (to ABT) and BB/S000453/1 (to GM and ABT and BDH).

## Supplementary Figures

**Figure S1.**
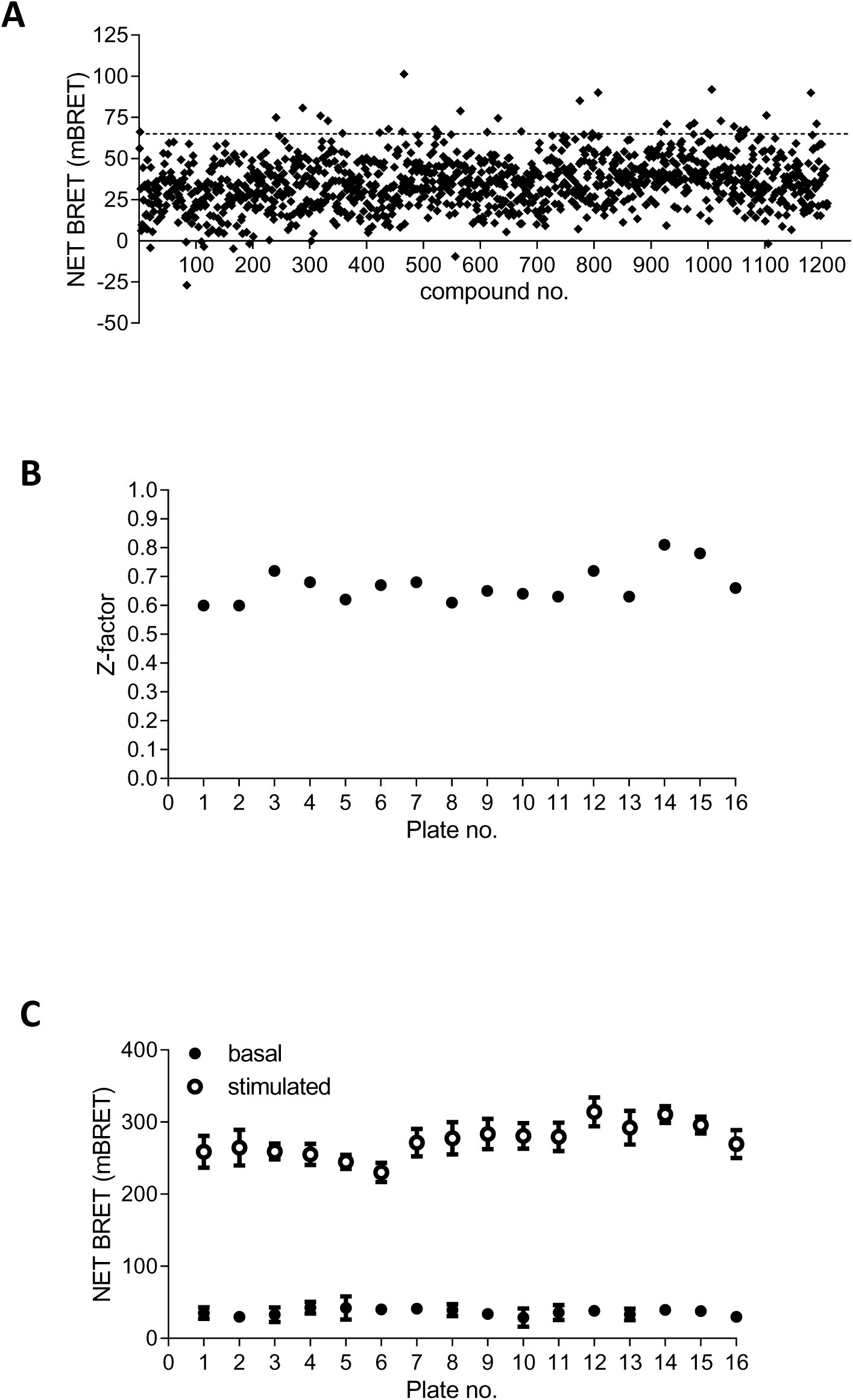
Development of a screening assay to identify novel hFFA2-DREADD activators. **A.** Screening of 1210 compounds related to sorbate at 100 μM in a hFFA2-DREADD-β-arrestin-2 interaction identified various potential hits in a single point assay. Data are presented as Net milliBRET. The dotted line represents mean + 3 S.D. **B.** The Z’ factor (ref 23) of assay robustness was calculated across individual 96 well microtitre plates of compounds screened in A. **C.** Signal to background across the assay format was assessed by spiking 4 wells of each plate with 100 μM sorbate (**open symbols**) as a positive control and comparing this with a vehicle control (**filled symbols**). Data are means +/- S.E.M.

**Figure S2.**
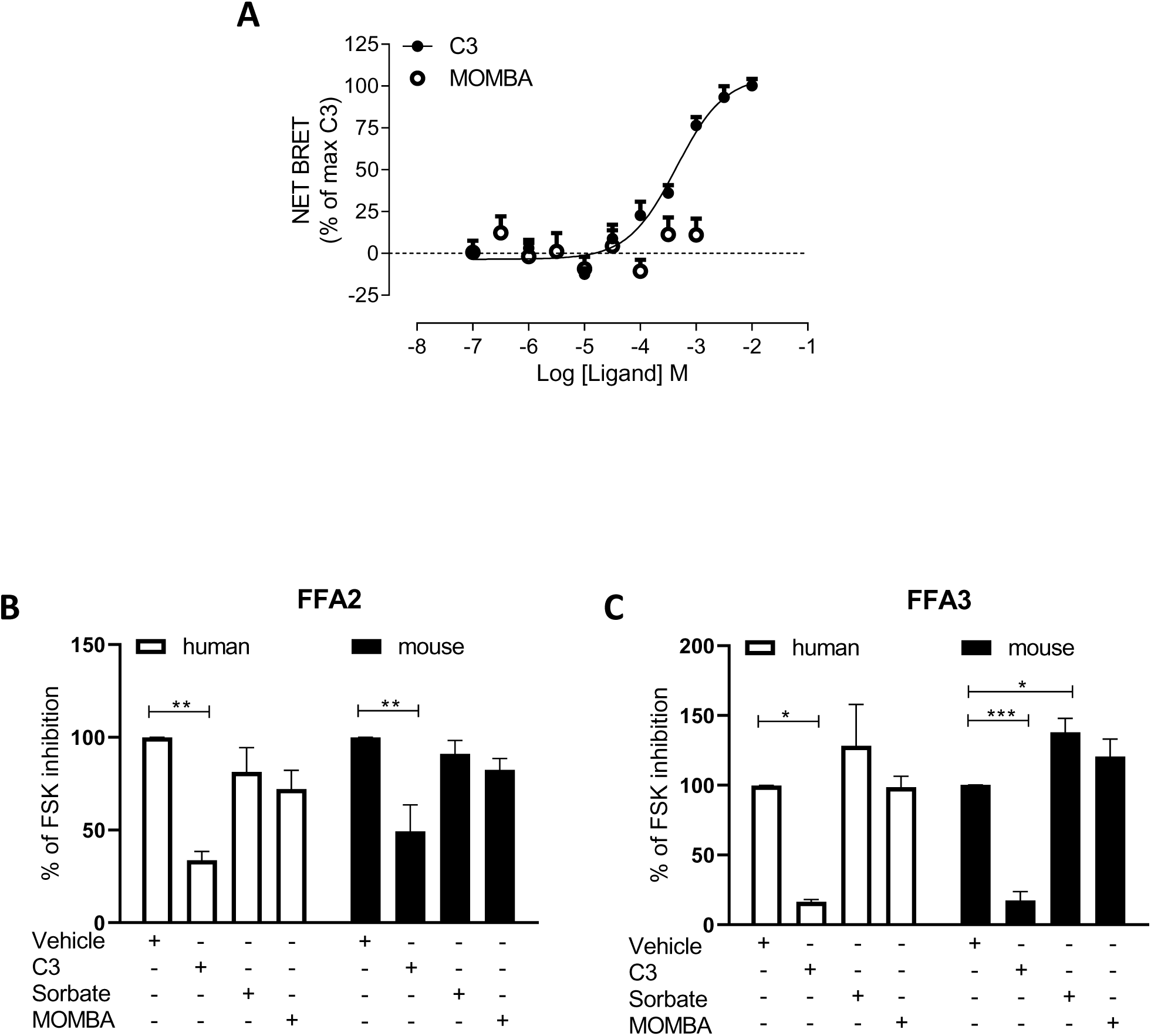
MOMBA is a selective agonist at hFFA2-DREADD. A. The ability of varying concentrations of C3 (**filled symbols**) or MOMBA (**open symbols**) to promote interactions between wild type human FFA2 and β-arrestin-2 was assessed in a BRET-based assay. Data are means +/- S.E.M. from 3 separate experiments. B. The ability of a single concentration of C3 (1 mM), sorbate (1 mM), and MOMBA (100 μM) to inhibit forskolin-stimulated levels of cAMP in HEK293 cells induced to express either human (**open bars**) or mouse (**filled bars**) FFA2 was compared to vehicle. Data are means +/_ S.E.M. n = 3, ** P < 0.01. C. Studies equivalent to B were performed in HEK293 cells induced to express either human (**open bars**) or mouse (**filled bars**) FFA3. Data are means +/_ S.E.M. n = 3, * P < 0.05, *** P < 0.001.

**Figure S3.**
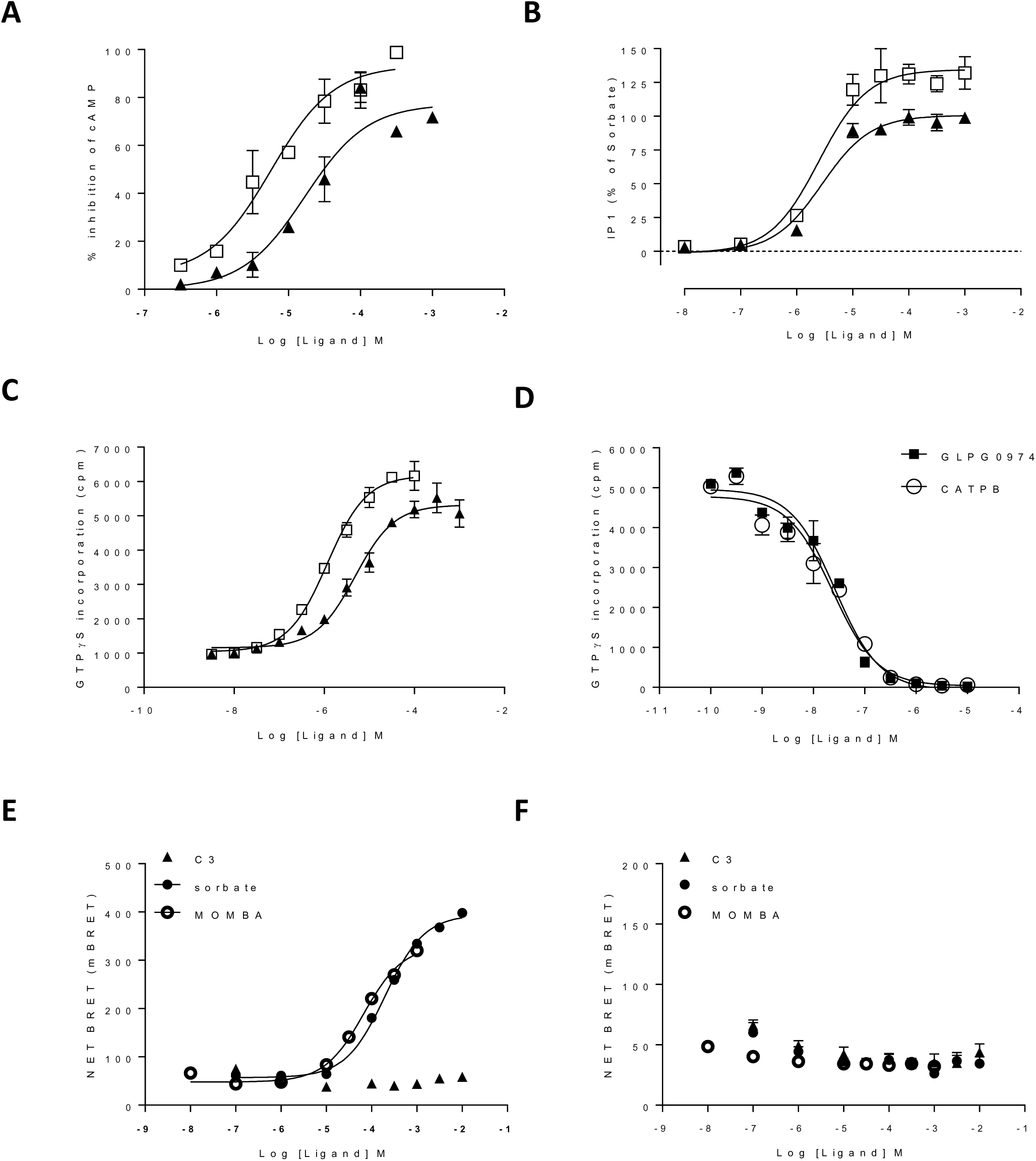
MOMBA is an effective and more potent agonist of hFFA2-DREADD than sorbate. A. The ability of varying concentrations of either MOMBA (**open symbols**) or sorbate (**closed symbols**) to inhibit forskolin-stimulated cAMP levels was assessed in cells induced to express hFFA2-DREADD. B. The ability of varying concentrations of either MOMBA (**open symbols**) or sorbate (**closed symbols**) to promote production of inositol monophosphates in cells induced to express hFFA2-DREADD is shown. C. Membrane preparations of cells induced to express hFFA2-DREADD were used to assess the ability of MOMBA (**open symbols**) or sorbate (**closed symbols**) to promote binding of [^35^S]GTPγS. D. The capacity of varying concentrations of either CATPB (**open symbols**) or GLPG0974 (**filled symbols**) to inhibit binding of [^35^S]GTPγS induced by an EC_80_ concentration of MOMBA was assessed. E. MOMBA and sorbate was unable to activate an orthosteric (Arg^180^Ala) binding site mutant of hFFA2-DREADD in a receptor-β-arrestin-2 interaction assay.

**Figure S4.**
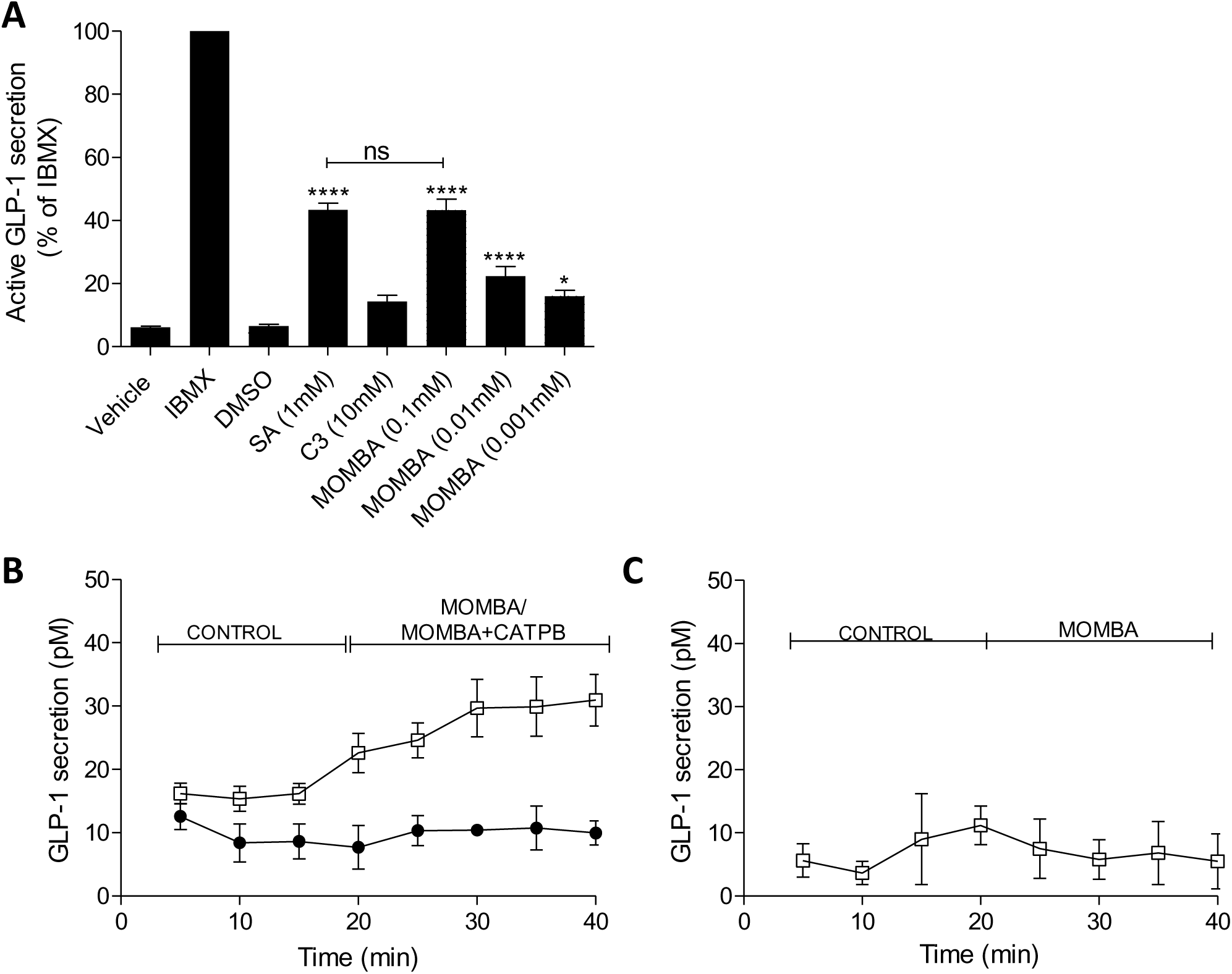
MOMBA promotes release of GLP-1 in colonic preparations from hFFA2-DREADD-HA expressing mice. A. Colonic crypt-containing preparations were isolated from hFFA2-DREADD-HA expressing mice. MOMBA promoted release of GLP-1 in a concentration-dependent fashion (* P < 0.05, **** P < 0.0001) whilst C3 did not. At maximally effective concentrations MOMBA was as effective as sorbate (SA) (ns = not significantly different). Isobutylmethylxanthine (IBMX) (100 μM) provided a positive control for release of GLP-1. Data represent means +/- S.E.M from 5 different preparations. B. MOMBA (**open symbols**) promoted sustained release of GLP-1 release from colonic tissue of hFFA2-DREADD-HA expressing mice and this was prevented by co-exposure to CATPB (**filled symbols**). Data represent means +/- S.E.M from 4 different preparations (*** P < 0.001). C. MOMBA (**open symbols**) did not promote release of GLP-1 from colonic tissue obtained from CRE-MINUS animals. Data represent means +/- S.E.M from 4 different preparations.

**Figure S5.**
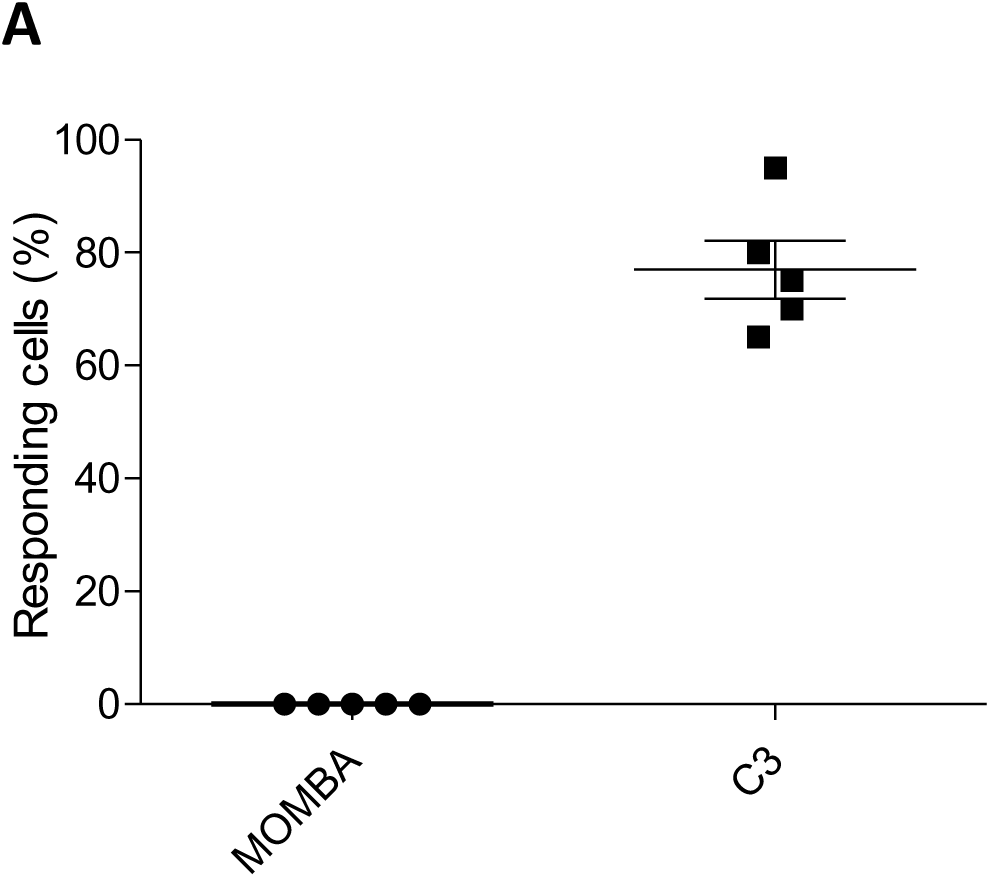
MOMBA is unable to stimulate increases in Ca^2+^ in DRG cells from mice lacking FFA2. Cell dispersed from DRGs isolated from CRE-MINUS mice were used to assess the ability of ligands to elevate Ca^2+^ as in Figure 6. No effect of MOMBA was recorded whilst C3 was effective in most of the cells tested.

**Supplementary Table 1.** MOMBA promotes Ca^2+^ elevation in subsets of DRG-derived cells in a FFA2 and G_q_/G_11_-dependent manner

|  | **MOMBA** | **MOMBA +**<br>**CATPB** | **MOMBA +**<br>**FR900359** | **MOMBA +**<br>**P.toxin** |
| --- | --- | --- | --- | --- |
| Animals (N) | 9 | 7 | 5 | 6 |
| Cells (n) | 103 | 68 | 63 | 74 |
| % activated<br>Mean +/-<br>S.E.M. | 41.4 +/- 10.0 | 2.6 +/- 1.7 | 3.4 +/- 1.9 | 39.2 +/- 4.9 |

**Supplementary Table 2.** C3 promotes Ca^2+^ elevation in subsets of DRG-derived cells in a non-FFA2 and G_i_-dependent manner

|  | **C3** | **C3 + CATPB** | **C3 + FR900359** | **C3 + P.toxin** |
| --- | --- | --- | --- | --- |
| Animals (N) | 7 | 5 | 4 | 5 |
| Cells (n) | 84 | 41 | 52 | 59 |
| % activated<br>Mean +/-<br>S.E.M. | 81.1 +/- 3.8 | 73.0 +/- 6.6 | 82.5 +/- 4.3 | 11.0 +/- 5.6 |

**Supplementary Table 3.**
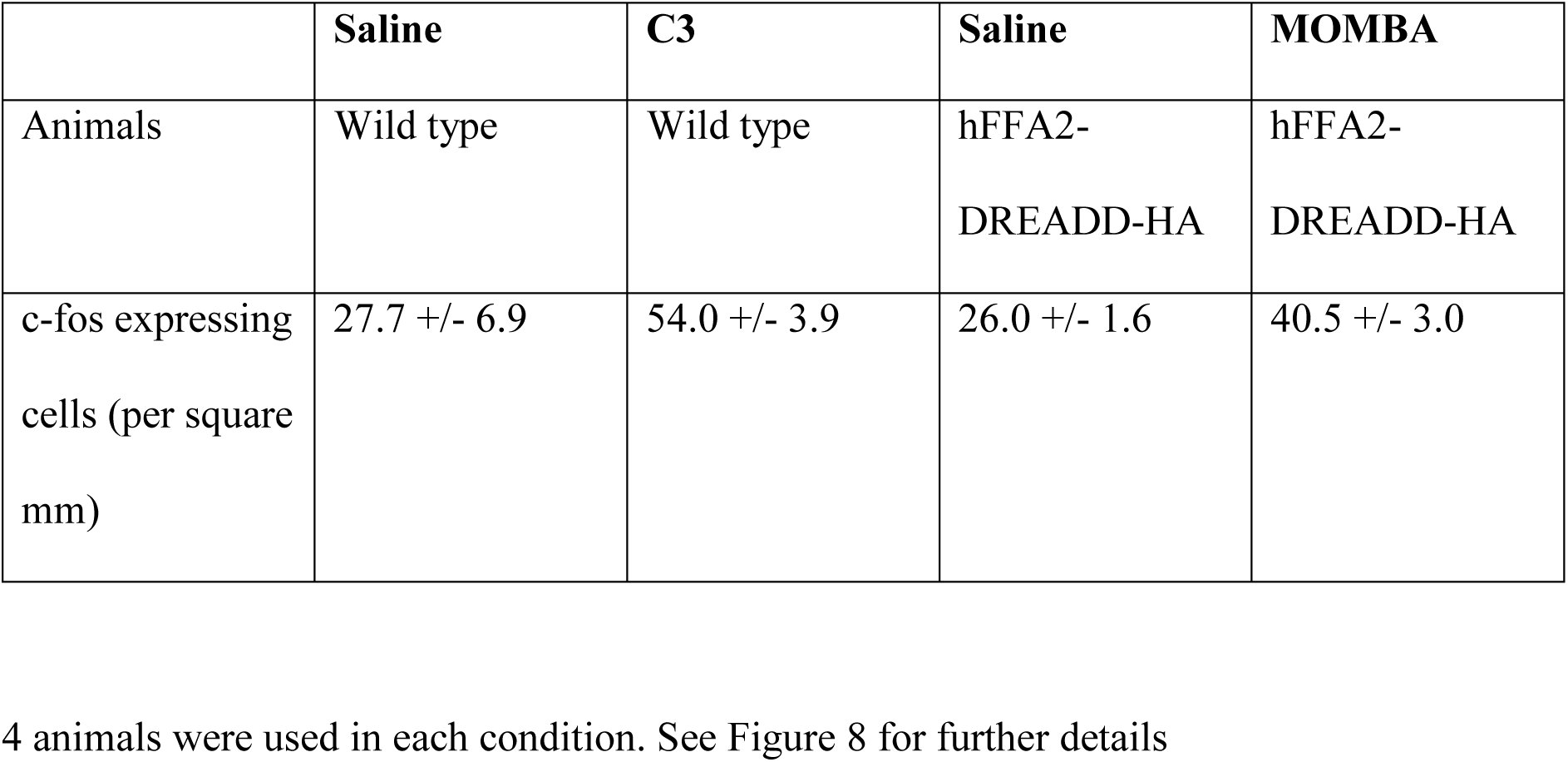
Activation of gut SCFA receptors promotes c-fos expression in the dorsal horn of the spinal cord

